# ReDisX: a Continuous Max Flow-based framework to redefine the diagnosis of diseases based on identified patterns of genomic signatures

**DOI:** 10.1101/2022.04.11.487592

**Authors:** Hiu Fung Yip, Debajyoti Chowdhury, Kexin Wang, Yujie Liu, Yao Gao, Liang Lan, Chaochao Zheng, Daogang Guan, Kei Fong Lam, Hailong Zhu, Xuecheng Tai, Aiping Lu

## Abstract

Diseases originate at the molecular-genetic layer, manifest through altered biochemical homeostasis, and develop symptoms later. Hence symptomatic diagnosis is inadequate to explain the underlying molecular-genetic abnormality and individual genomic disparities. The current trends include molecular-genetic information relying on algorithms to recognize the disease subtypes through gene expressions. Despite their disposition toward disease-specific heterogeneity and cross-disease homogeneity, a gap still exists to describe the extent of homogeneity within the heterogeneous subpopulation of different diseases. They are limited to obtaining the holistic sense of the whole genome-based diagnosis resulting in inaccurate diagnosis and subsequent management.

To fill those gaps, we proposed ReDisX framework, a scalable machine learning algorithm that uniquely classifies patients based on their genomic signatures. It was deployed to re-categorizes the patients with rheumatoid arthritis and coronary artery disease. It reveals heterogeneous subpopulations within a disease and homogenous subpopulations across different diseases. Besides, it identifies *GZMB* as a subpopulation-differentiation marker that plausibly serves as a prominent indicator for *GZMB*-targeted drug repurposing.

The ReDisX framework offers a novel strategy to redefine disease diagnosis through characterizing personalized genomic signatures. It may rejuvenate the landscape of precision and personalized diagnosis, and a clue to drug repurposing.

## Introduction

Transforming the conventional diagnosis strategy for diseases is emerging for ensuring a healthier lifespan (Zhao et al., 2020) and for strengthening drug repurposing (Pushpakom et al., 2019). In the modern era, the practices for diagnosing diseases have been progressively oriented towards precision and personalized, relying on a molecular genetic basis, especially gene expression-based (Aure et al., 2017; He et al., 2021). Since the conventional diagnosis of diseases often remains insufficient in explaining heterogeneity within a disease and the homogeneity between multiple diseases (Humby et al., 2019; Khera & Kathiresan, 2017). Transcriptomic studies demonstrated that the molecular heterogeneity within one disease would be highly divergent, for example, colon cancer (Marisa et al., 2013)(Marisa et al., 2013), breast cancer (Higgins & Baselga, 2011; Yu et al., 2019)(Higgins & Baselga, 2011; Yu et al., 2019), rheumatoid arthritis (RA) (Humby et al., 2019)(Humby et al., 2019), and coronary artery disease (CAD) (Khera & Kathiresan, 2017)(Khera & Kathiresan, 2017). Alongside the heterogeneity within a disease, the homogeneity among different diseases is also a critical aspect to study. For example, RA and CAD have shared a similar inflammatory pathway (Hansson, 2005; Lee & Weinblatt, 2001; Lee et al., 2019)(Hansson, 2005; D. M. Lee & Weinblatt, 2001; T. H. Lee, Song, Choi, Seok, & Jung, 2019). Niu et al. discovered four shared canonical pathways, three shared networks, and three upstream regulators-driven inflammatory activations across RA and CAD (Niu et al., 2014)(Niu et al., 2014). Offering symptomatic managements without considering the deeper knowledge of underpinning heterogeneity may result in treatment failure and/or resistance to the drugs (Lim & Ma, 2019; Ouboussad et al., 2019)(Lim & Ma, 2019; Ouboussad, Burska, Melville, & Buch, 2019; Udalova, Mantovani, & Feldmann, 2016).

It certainly embarks us to investigate the initialization of the diseases. Any clinicopathological condition or disease is typically originated at the gene level, and the associated altered physiological states and biochemical balances are manifested as phenotypes and then symptoms (Gao et al., 2018; Larsen & Minna, 2011). Hence, the clinical symptoms used to reflect the imbalance of physiological homeostasis. It usually does not comprise the actual underlying disparities at the point of origin which is at the molecular-genetic layer (Gao et al., 2018; Larsen & Minna, 2011). Therefore, defining the diseases based on clinical symptoms may not encompass the whole underlying disparities at the molecular-genetic level. It is even more unclear when the same gene critically plays a role in manifesting two different disease phenotypes (Afewerky, 2020; Grizzanti et al., 2019; Niu et al., 2014), which may lead to an erroneous diagnosis and could end up with receiving decisive treatments (Zhao et al., 2020). It also hinders the possibility of repurposing some drugs (Pushpakom et al., 2019). For instance, Aspirin is typically used for managing analgesia, and it was successfully repurposed to manage Colorectal cancer as both share the same genetic causes (Pushpakom et al., 2019).

Enabling precision diagnosis and personalized medicine using unprecedented molecular-level data is playing a major role in modern medicine. The deep multi-view learning approach demonstrated the power of integrated multi-omics data to identify potential biomarkers for specific disease types (T. Wang et al., 2021)(Wang et al., 2021). Several other studies also strengthened the engagement of multi-omics data-driven classifications of the in-disease expression differentiation that elucidates underlying pathological diversities (Dash et al., 2019; Li et al., 2015)(Dash, Shakyawar, Sharma, & Kaushik, 2019; Li et al., 2015). Despite having many advanced tools, most of the available frameworks are merely not adequate to discover the disease heterogeneity. They mostly rely on clustering algorithms those group the patients by unsupervised learning upon discovering similar gene expression profiles and then annotating their pathological properties (Aure et al., 2017; He et al., 2021). Clustering approaches of both types, deep learning (DL) and machine learning (ML) (Li et al., 2015) based algorithms have demonstrated their respective competence in investigating the heterogeneity within disease and homogeneity between multiple diseases. However, many existing ML approaches for clustering were slightly out-of-context here as they only focus on heterogeneity within one disease and do not encompass cross disease similarity within the heterogonous subpopulation. They also fail to specify the optimal number of clusters to be distinctly discovered.

Here, we have proposed ReDisX (Redefining the Disease X) framework to address all those ambiguities described above. It is an integrated ML clustering algorithm relying on a Continuous Max Flow (CMF) model (Wei et al., 2017; Yin & Tai, 2018) that uses a hierarchical clustering approach to pre-label the input patient data and then the CMF model to predict the patient labels uniquely. ReDisX re-categorizes the diseases based on the heterogeneity of individual gene expression profiles of the patients. Then, it evaluates the cross-disease similarity in expression level to discover whether two different diseases share the same or similar gene expression profiles. This paper considers CAD and RA as our diseases of interest-based on prior research experience (Niu et al., 2014). The ReDisX is self-sufficient in evaluating the optimal number of clusters.

Our results revealed an enhanced differentiation capacity compared to the conventional diagnosis (Figure 2, 3). ReDisX showed distinct differentiation ability across the diversified datasets (Figure 2, 3). It showed that a subpopulation (n=12) of CAD patients exhibits higher gene expression similarity and functional enrichment similarity to the RA patients (Figure 4). It clustered the similar gene expression profiles of the subpopulation of CAD and RA consecutively to classify the heterogeneity within a disease. Interestingly, a hub-gene was discovered within the CAD and RA subpopulation related to the drug discovery targeting *GZMB* (Figure 4) (Joehanes et al., 2013). Our analyses have endorsed *GMZB* as a potential target for developing drugs for those specific subpopulations of CAD and RA.

ReDisX may offer a clue to repurpose any potential drug candidates discovered for CAD that can be validated for RA and vice versa. It reinforces a novel ML approach to redefine the existing two diseases into a total of eight (three within RA and five within CAD) distinct categories based on gene expression signatures. This data-driven novel approach offers an enhanced resolution in introducing better precision and a personalized diagnosis strategy.

## RESULTS

We have developed a novel framework, ReDisX, an integrated ML algorithm relying on the CMF model. It intends to re-categorize the patient populations and their clinical conditions based on specific gene expression signatures (Figure 1A). Our study considered the RA and CAD patient’s data as we had prior expertise in that regime (Niu et al., 2014).

**Figure 1:**
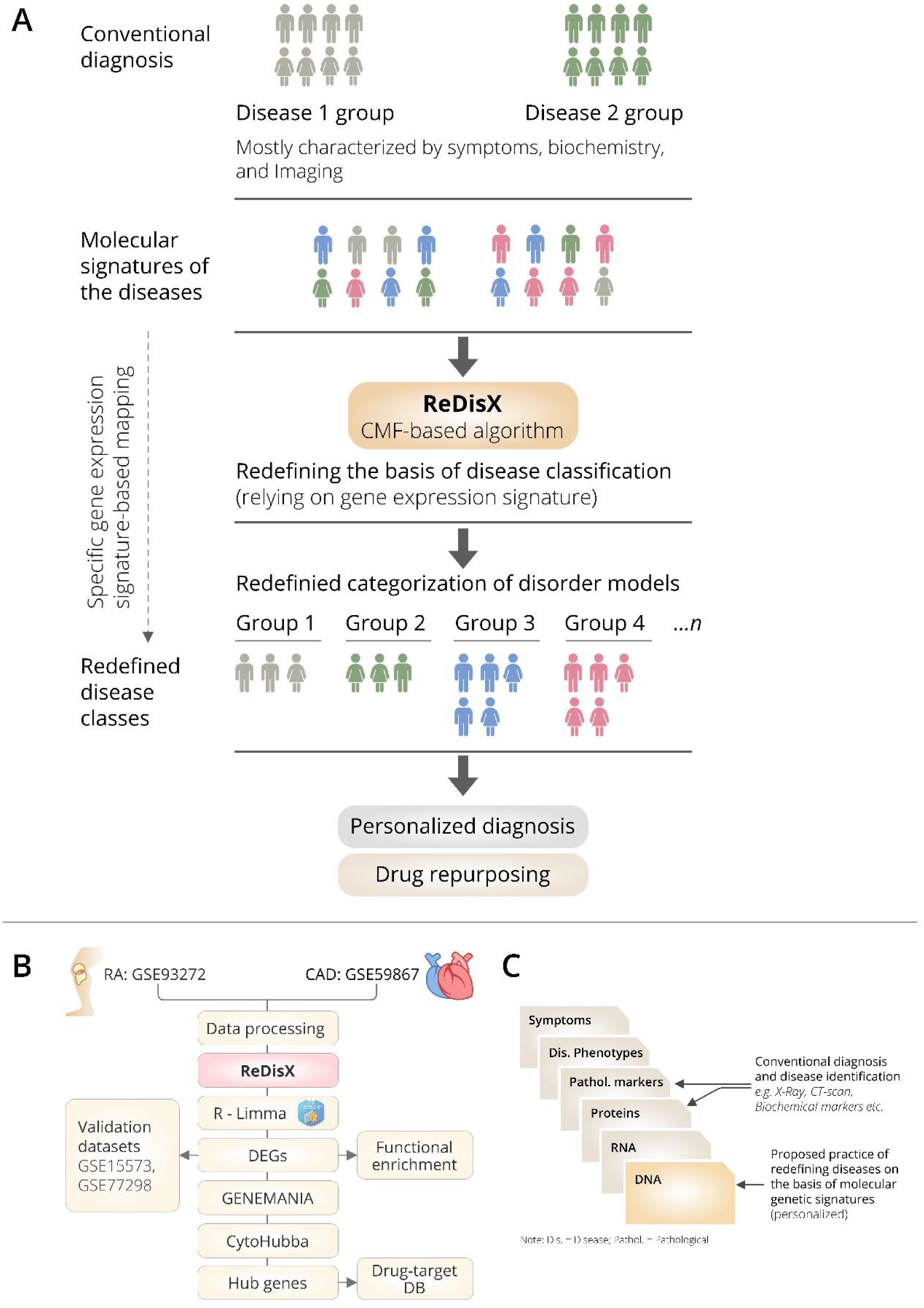
The schematic illustration of the ReDisX framework and its functional features. (A) ReDisX-based approach to redefine the disease diagnosis compared to the conventional diagnosis, (B) functional pipeline for the ReDisX-based analyses for RA and CAD datasets, (C) strategic significance of ReDisX framework over the conventional diagnosis procedure.

The ReDisX considers the gene expression data as an input to execute further analyses to identify the core of the core signature genes, which may correspond to the unique features of a clinical condition for an individual (Figure 1B). Our study indicated a distinctive classification among the patients diagnosed with RA and CAD. It also discovers a subpopulation of patients sharing similar characteristics despite being heterogeneous diagnosed as a separate disease. Ultimately, ReDisX reinforces a fresh data-driven perception to reconsider the basis of diagnosing the clinical conditions, relying on precision focus at the molecular genetic level (Figure 1C). It showed a clue to navigate a druggable target for a disease where a formula originally discovered for other diseases shares a strength to be validated.

### ReDisX redefines RA patients through acquiring personalized and precise molecular-genetic information

ReDisX serves as a core framework to extract the underlying transcriptional heterogeneity of the RA patient population. To identify the optimal number of heterogeneous sub-populations in RA patients, we employed the ReDisX cluster evaluation in human whole blood transcriptome data (GSE93272), which measure the mRNA expression (Tasaki et al., 2018). ReDisX adopted the optimal number (k=3) of heterogeneous sub-populations in the RA patients and grouped the high similarity patients into the same cluster (Figure 2D).

**Figure 2:**
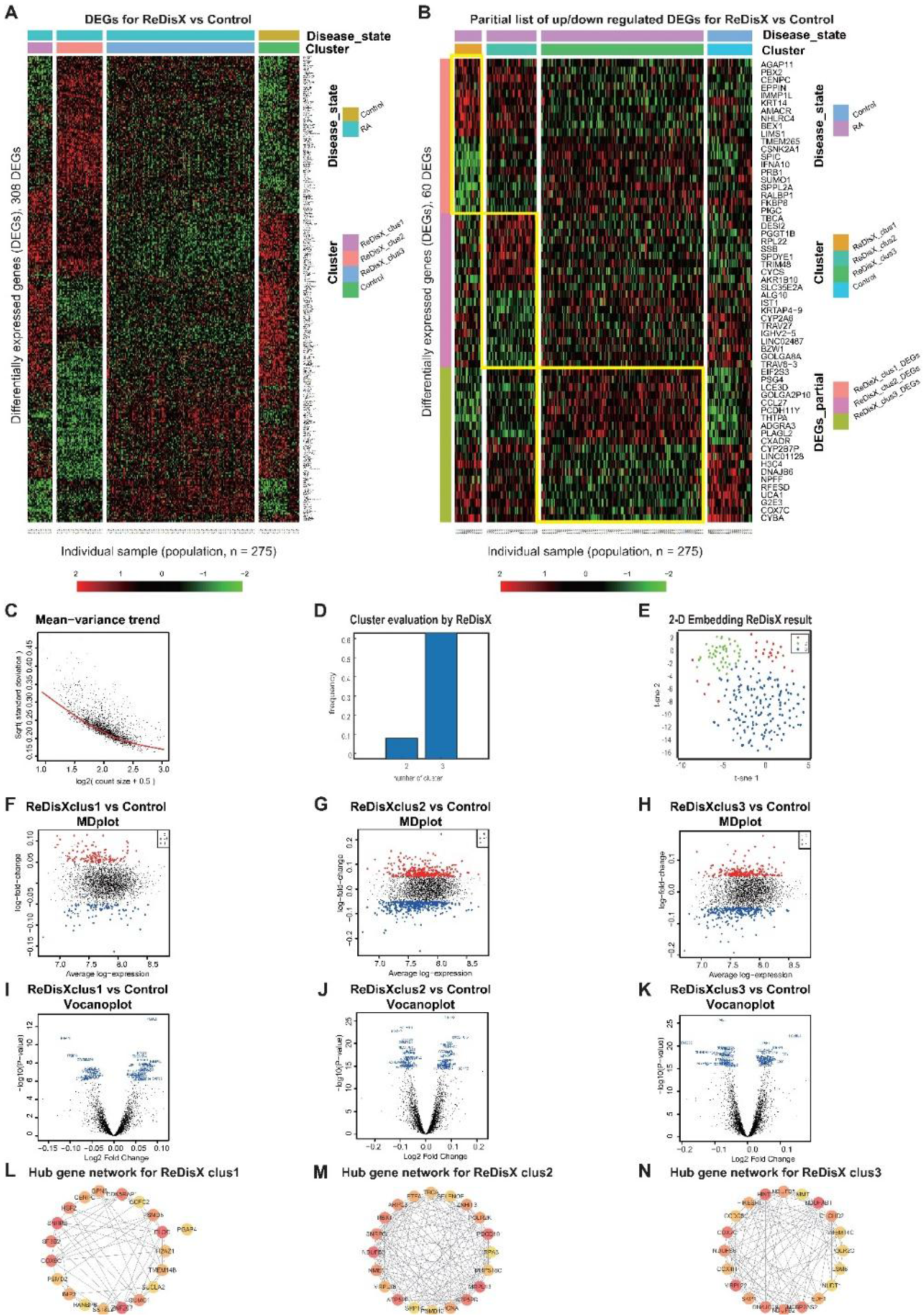
ReDisX redefines the RA patients through acquiring personalized and précised molecular-genetic information. (A) Heatmap for relative expression in up/downregulating DEGs identified in GSE93272 (p-value <0.05, log fold change>0.05, non-overlapping DEGs with other subpopulations). (B) Randomly selected 10 up-regulating genes and 10 downregulating genes corresponding to each ReDisX-based cluster in GSE93272 (p-value <0.05, log fold change>0.05, non-overlapping DEGs with other subpopulation). (C) Mean-variance trend after processed voom in GSE93272. (D) Optimal subpopulations were discovered from the total number of 232 RA patients in GSE93272 by ReDisX. (E) PCA and t-SNE visualization of the total number of 232 RA patients in GSE93272. (F) Mean-Difference (MD) plots for the ReDisX-based cluster 1 of RA patients to the healthy controls [label 1 (red dot) represents the up-regulated genes in the upper panel, the label −1 (blue dot) represents the down-regulated genes at the lower panel]. (G) MD plots for ReDisX-based cluster 2 of the RA patients to the healthy controls [label 1 (red dot) represents the up-regulated genes, and the label −1 (blue dot) represents the down-regulated genes]. (H) MD plots for ReDisX-based cluster 3 to the healthy controls [label 1 (red dot) represents the up-regulated genes, label −1 (blue dot) represents the down-regulated genes]. (I) Volcano plot for ReDisX-based cluster 1 compared to the healthy controls with highlighting top 50 log2 fold change genes in blue. (J) Volcano plot for ReDisX-based cluster 2 compared to healthy controls with highlighting top 50 log2 fold change genes in blue. (K) Volcano plot for ReDisX-based cluster 3 compared to healthy controls with highlighting top 50 log2 fold change genes in blue. (L-N) cytoHubba-identified hub gene analysis to the GeneMANIA network for ReDisX-based clusters using the MCC ranking algorithm. The hub genes for cluster 1 (L), cluster 2 (M), the cluster 3 (N) were shown.

As per ReDisX-based patient labelling, differential expressions (DE) analyses were performed on each sub-population using Limma in R (Ritchie et al., 2015). After excluding the overlapping differentially expressed genes (DEGs) among the subpopulations, a total number of 308 genes were found to be differentially expressed (p-value <0.05, log fold change>0.05) (Figure 2A). To provide a higher resolution in the co-expression patterns based on the ReDisX discovery, we randomly selected 10 up-regulated genes and 10 downregulated genes for each ReDisX-based cluster. It showed 60 genes in rows and 275 samples in columns (Figure 2B).

Further, the network analyses for those DEGs were conducted using GeneMANIA (Warde-Farley et al., 2010), a Cytoscape function (Shannon et al., 2003). It is constructed using six established co-expression analysis tools (Agnelli et al., 2007; Agnelli et al., 2009; Brodmerkel et al., 2019; B. Zhang et al., 2020; H. Zhang et al., 2020). Next, we estimated the Maximal Clique Centrality (MCC) score that predicts essential nodes within the biological networks in cytoHubba (Chin et al., 2014). Higher MCC scores indicated the more critical hub genes. Then, the top 20 hub genes were selected from each subpopulation of patients (Figure 2L-2N).

The ReDisX framework identified the following hub genes. *SUMO1, PSMD6, SNRPB, H2AZ1, RANBP6, SUCLA2, GPN3, PGAP4, CENPC, PSMD2, ZNF207, GCFC2, SS18L2, COX6C, ELOC, TMEM14B, HSF2, IMP3, CDK5RAP1* for the cluster 1 (Figure 2L), *ATP5PB, MRPS18C, MRPL36, SRP14, PSMD10, NDUFB3, TBCA, ZNHIT3, NME1, RPA3, RBX1, MRPL13, SELENOF, PDCD10, PCNA, ATP5PF, ETFA, SNRPG, ARPC3, POLR2K* for the cluster 2 (Figure 2M), and *COX4I1, SKP1, DNAJC19, NDUFB2, NCBP2AS2, EDF1, IMMT, POLR2G, COX7C, NUDT1, HIKESHI, CCDC51, NDUFB5, MRPL22, HINT1, TMEM14C, LSM6, CHCHD2, NDUFS6, NDUFAB1* for the cluster 3 (Figure 2M) were identified and their corresponding networks were constructed (Figure 2L-2N). The detailed Cytoscape networks are available in supplementary 2.

Visualization of the gene expression profiles for RA patients was reduced to 50 dimensions by principal component analyses (PCA) (Hotelling, 1933; Pearson, 1901), then further reduced to 2 dimensions by t-distributed stochastic neighbor embedding (t-SNE) (van der Maaten & Hinton, 2008). It returned a final two dimensions plot consisting of 232 patients’ data (Figure 2E). Finally, quality control of the data was ensured using a mean-variance trend (Figure 2C). To support it, Mean Difference (MD) plots were plotted for each ReDisX-based cluster to the healthy controls (Figure 2F-2H). The volcano plots for each ReDisX-based cluster compared to the healthy controls also highlighted the top 50 log2 fold change genes in blue (Figure 2I-2K).

### ReDisX distinguishes the heterogeneous subpopulation among the CAD patients

Our results also showed the efficiency of ReDisX to extract the underlying transcriptional heterogeneity of the CAD patients. The analyses pipeline was similar to the previous section on analyzing RA patients. To identify the optimal number of heterogenous subpopulations in CAD patients, we employed the ReDisX cluster evaluation in human whole blood transcriptome data (GSE59867) that measures the mRNA expression (Maciejak et al., 2015). ReDisX adopted the optimal number (k=5) for heterogenous subpopulations in the CAD patients and grouped the highly similar patients into the same cluster (Figure 3D).

**Figure 3:**
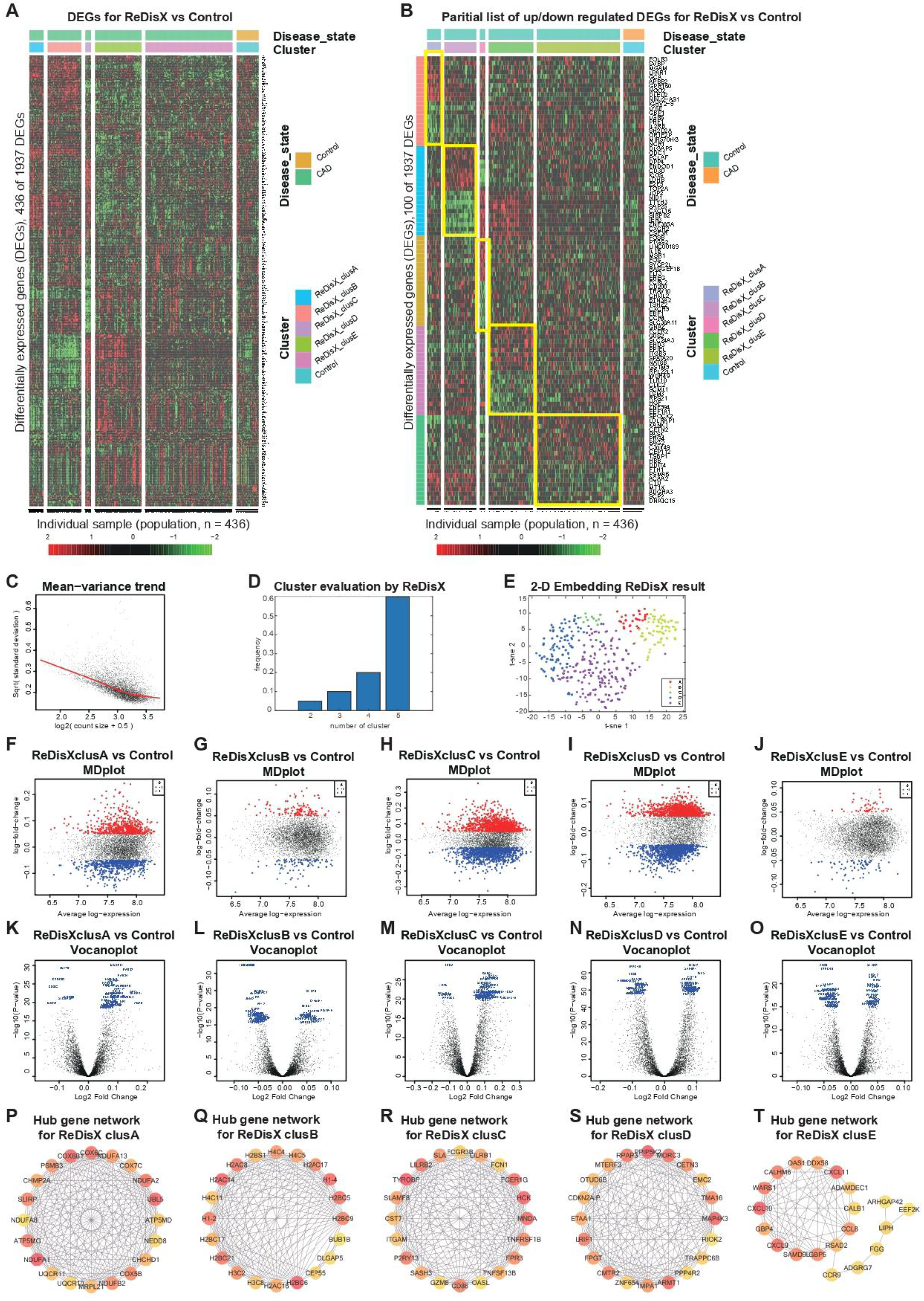
ReDisX distinguishes the heterogeneous subpopulation among CAD patients. (A) Heatmap for relative expression in up/downregulating DEGs identified in GSE59867 (p-value <0.005, non-overlapping DEGs with other subpopulations). (B) Randomly selected 10 up-regulating genes and 10 downregulating genes correspond to each ReDisX-based cluster in GSE59867 (p-value <0.005, non-overlapping DEGs with other subpopulations). (C) Mean-variance trend after processed voom in GSE59867. (D) Optimal subpopulations were discovered from the total number of 390 CAD patients in GSE59867 by ReDisX. (E) PCA and t-SNE visualization of the total number of 390 patients in GSE59867. (F-J) MD plots for ReDisX-based clusters. The ReDisX-based cluster A (F), cluster B (G), cluster C (H), cluster D (I) and the cluster E (J) of the CAD patients to the healthy controls were shown to have up/down-regulated genes (red dot represent the upregulated genes in the upper panel and the blue dot represents the downregulated genes in the lower panel). (K-O) Volcano plots for the ReDisX-based clusters compared to the healthy controls were highlighted using the top 50 log2 fold change genes in blue. ReDisX-based cluster A (K), cluster B (L), cluster C (M), cluster D (N), and cluster E (O) were shown to have a comparative difference of those top 50 log2 fold change genes to the healthy controls. (P-T) cytoHubba-identified hub gene analysis to the GeneMANIA network for ReDisX-based clusters using the MCC ranking algorithm. The hub genes for cluster A (P), cluster B (Q), cluster C (R), cluster D (S), and cluster E (T) were shown.

Then, DE analyses were performed on each subpopulation within the ReDisX-based patient labelling using Limma in R (Ritchie et al., 2015). After excluding the overlapping DEGs among the subpopulations, a total number of 436 out of 1937 genes were found to be differentially expressed (p-value <0.005) (Figure 3A). To provide a higher resolution in the co-expression patterns based on the ReDisX discovery, we randomly selected 10 up-regulated genes and 10 downregulated genes for each ReDisX-based cluster. It showed 100 genes in rows and 436 samples in columns (Figure 3B).

Furthermore, the network analyses were conducted for further filtered DEGs (p-value <0.05, log fold change>0.05) were conducted using GeneMANIA (Warde-Farley et al., 2010). We estimated the MCC score to predict the critical nodes within the biological networks (Chin et al., 2014). Higher MCC scores indicated the more important hub genes here as well. Then, the top 20 hub genes were selected from each subpopulation of patients (Figure 3P-3T) for further analyses.

The ReDisX framework identified the following hub genes. *MRPL21, UQCR10, NDUFB2, COX5B, CHCHD1, NEDD8, COX7C, CHMP2A, NDUFA8, SLIRP, PSMB3, NDUFA1, UQCR11, COX6C, COX6B1, ATP5MG, UBL5, NDUFA2, NDUFA13, ATP5MD* for the cluster A (Figure 3P), *H3C2, H3C8, H2AC16, H2BC6, CEP55, DLGAP5, H2BC9, H2BC5, H1-4, H2BS1, BUB1B, H1-2, H2AC8, H2BC21, H2AC17, H4C5, H4C4, H2AC14, H4C11, H2BC17* for the cluster B (Figure 3Q), and *TGAM, TNFSF13B, CD86, FPR3, P2RY13, GZMB, OASL, TNFRSF1B, SASH3, HCK, FCER1G, FCN1, CST7, LILRB2, LILRB1, TYROBP, MNDA, FCGR3B, SLAMF8, SLA* for the cluster C (Figure 3R), *PPP4R2, TRAPPC6B, RIOK2, ZNF654, IMPA1, RPAP3, EMC2, LRIF1, FPGT, TMA16, MORC3, PPIP5K2, OTUD6B, ARMT1, MAP4K3, ETAA1, CDKN2AIP, MTERF3, CETN3, CMTR2* for cluster D (Figure 3S), *LIPH, ADGRG7, CXCL10, CXCL9, SAMD9L, CALHM6, ARHGAP42, DDX58, GBP5, EEF2K, WARS1, RSAD2, CCL8, FGG, ADAMDEC1, CXCL11, GBP4, CCR9, CALB1, OAS1* for cluster E (Figure 3T) were identified and their corresponding networks were constructed (Figure 3P-3T). The detailed Cytoscape networks are available in supplementary 1.

Visualization of the gene expression profiles for CAD patients was reduced to 50 dimensions by PCA (Hotelling, 1933; Pearson, 1901), then further reduced to 2 dimensions by t-SNE (van der Maaten & Hinton, 2008). It returned a final two dimensions plot consisting of 436 patients’ data (Figure 3E).

Finally, quality control of the data was ensured using a mean-variance trend (Figure 3C). MD plots were plotted for each ReDisX-based cluster against the healthy controls (Figure 3F-3J). Moreover, the volcano plots for each ReDisX-based cluster compared to the healthy controls were also shown, highlighting the top 50 log2 fold change genes in blue (Figure 3K-3O).

### ReDisX discovers the cross-subpopulation homogeneity among the RA and CAD patients embarking on the redefinition of their diagnosis at molecular-level

ReDisX revealed the cross-subpopulation homogeneity among the CAD and RA patients by analyzing their transcriptomic profiles (Figure 4). The extent of the homogeneity was assessed in terms of the intersection of DEGs (Figure 4A), functional enrichment analysis (Figure 4B-4F) (Chen et al., 2013; Kuleshov et al., 2016) and the drug bank (Figure 4G) (Wishart et al., 2006) across the sub-population of CAD and RA.

**Figure 4:**
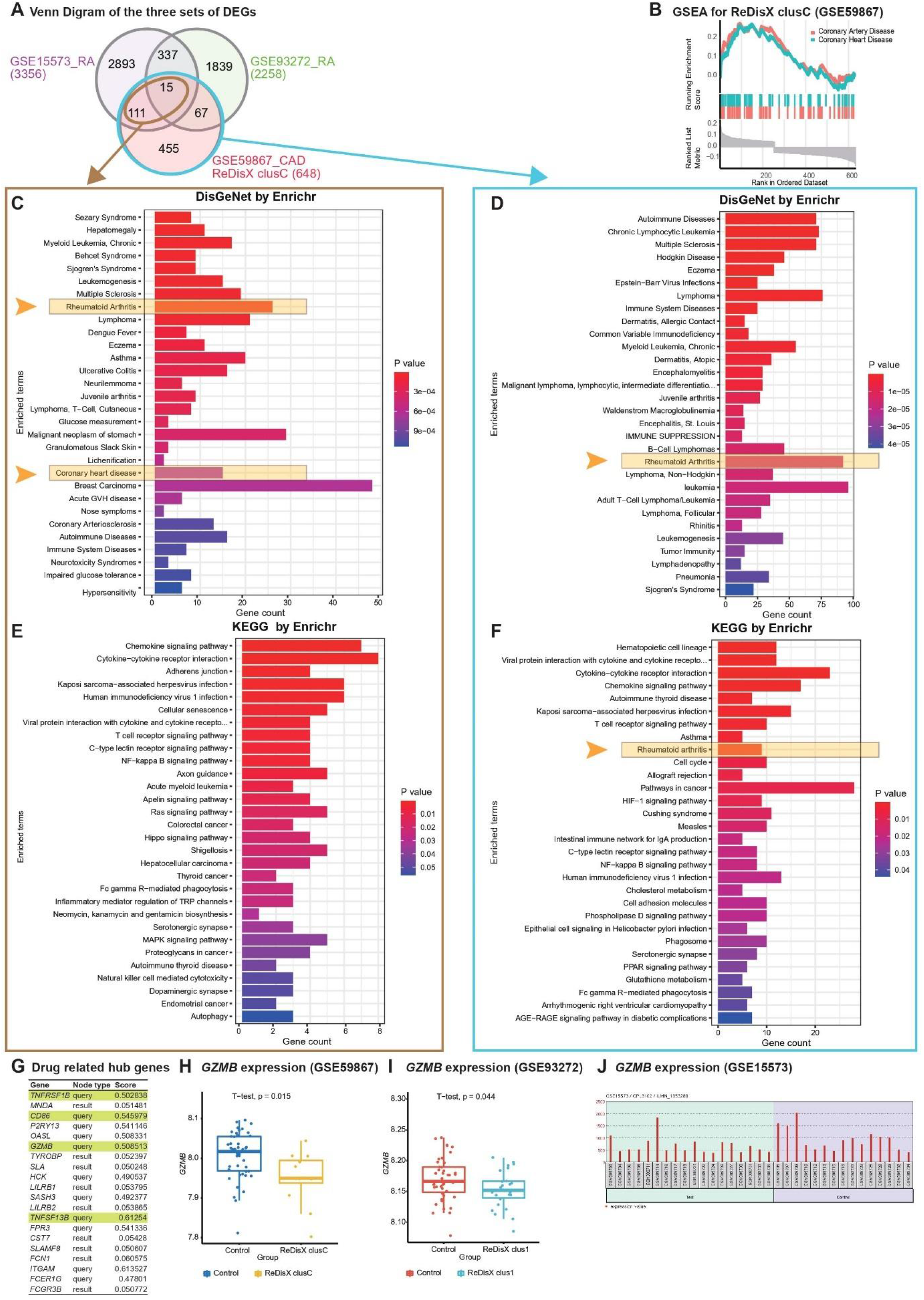
ReDisX-based discovery of the subpopulation homogeneity across the RA and CAD patients. (A) Venn Diagram of three sets of DEGs, GSE15573 (RA), GSE93272 (RA), GSE59867 (CAD) cluster C. (B) GSEA of ReDisX-based cluster C in GSE59867. (C) Disease Ontology (DO) enrichment using DisGeNet for the intersecting DEGs of GSE59867 (CAD) cluster C and GSE15573 using Enrichr. (D) DO enrichment of GSE59867 (CAD) cluster C DEGs. (E) Pathway enrichment for the intersecting DEGs of GSE59867 (CAD) cluster C and GSE15573 by KEGG. (F) Pathway enrichment for GSE59867 (CAD) cluster C DEGs by KEGG. (G) The list of drug-gene target related identified hub genes from the cluster C of GSE59867 (CAD). (H) Boxplot of *GZMB* expression in GSE59867, Control (blue) and ReDisX-based cluster C (yellow). (I) Boxplot of *GZMB* expression in GSE93272, Control (red) and ReDisX-based cluster C (blue). (J) The expression value of *GZMB* within the patient and healthy controls from GSE15573.

ReDisX-based CAD cluster C (12 patients) of GSE59867 was discovered to be homogeneous to RA patients. Based on ReDisX-based patient labelling, 648 DEGs were identified from the GSE59867 CAD cluster C (Figure 4A). To validate the homogeneity of GSE59867 cluster C to RA, we employed two publicly available validation datasets for RA, GSE15573 and GSE93272 retrieved from whole blood transcriptome. Then, GEO2R (Barrett et al., 2013) was applied to identify the DEGs with default parameters (p-value <0.05). A total of 3356 and 2258 DEGs were identified from GSE15573 and GSE93272, respectively (Figure 4A). Moreover, a total number of 126 genes were found at the intersection of GSE59867 CAD cluster C and GSE15573 RA. It sparked a notion further to investigate the homogeneity between CAD subpopulation and RA patients, especially to identify some clues about more precision yet personalized diagnosis and drug repurposing. Additional GO analysis is available in (supplementary 2.1). Then, we conducted Gene Set Enrichment Analysis (GSEA) using clusterProfiler (T. Wu et al., 2021) to ensure that the ReDisX framework does not lose any essential molecular functions associated with the identified DEGs from the subpopulation of cluster C of CAD patients (Figure 4B).

Further to examine the hypothesis above, a total number of 126 intersecting DEGs of GSE59867 CAD cluster C and GSE15573 RA were analyzed for their functional enrichment using Enrichr (Chen et al., 2013; Kuleshov et al., 2016) and also with two other databases, DisGeNet (Piñero et al., 2017) for identifying enrichment of diseases, and KEGG (Kanehisa & Goto, 2000) for common pathways. In the DisGeNet enrichment analysis of 126 intersecting DEGs, RA and coronary heart disease, which implies a broader term for CAD, were observed in the top 30 enriched Disease Ontology (DO) terms. Also, in the KEGG analysis of 126 intersecting DEGs, two essential inflammatory pathways, Chemokine signalling, and Cytokines-Cytokines receptor interaction were observed in the top 2 enriched KEGG terms. Additional GO analysis is available in (supplementary 2.2). Similar validations were also conducted for another RA dataset, GSE77298 from synovial biopsies tissue (supplementary 3).

In parallel to the intersecting DEGs of GSE59867 CAD cluster C and GSE15573 RA (Figure 4A), the functional enrichment of a total number of 648 DEGs of GSE59867 cluster C was analyzed using Enrichr (Chen et al., 2013; Kuleshov et al., 2016), with two databases, DisGeNet (Piñero et al., 2017) and KEGG (Kanehisa & Goto, 2000). In the DisGeNet enrichment analysis of those DEGs, we observed that RA was in the top 30 enriched DO terms. Also, In the KEGG enrichment analysis of those DEGs, we observed that RA, Chemokine signalling, and Cytokines-Cytokines receptor interaction were in the top 10 enriched KEGG terms.

To identify whether any distinguished subpopulation shares any drug-related targets and/or pathways, we employed the drug bank (Wishart et al., 2006) to retrieve the drug-gene target database to analyze the drug-related hub genes of cluster C of GSE59867 CAD. Of them, a total number of 4 genes (*TNFRSF1B, CD86, GZMB and TNFSF13B*) were identified as the drug-related hub genes (Figure 4G, highlighted in light green). Our analysis indicated that one of the drug-related hub genes, *GZMB* was under-expressed in the ReDisX-based cluster 1 of the RA patients and the ReDisX-based cluster C of the CAD patients. The gene, *GZMB* encodes the granzyme B, which is secreted by natural killer cells and cytotoxic T-lymphocytes to induce inflammatory reactions by processing cytokines and imparting into chronic inflammations, including RA (C.-X. Bao et al., 2018), and in cardiovascular diseases (Joehanes et al., 2013; Travers et al., 2016). It plausibly indicated a notion of homogeneity discovered across the subpopulation of RA and CAD patients. Our results also indicated that heterogeneous subpopulations in both RA (p-value=0.044) and CAD (p-value=0.015) were under-expressed compared to their controls (Figure 4H, 4I). The analysis was validated using GEO2R (Barrett et al., 2013) differential expression analysis in GSE15573 (Figure 4J).

Further, the STITCH analysis indicated a strong inter-relationship among the total identified 40 hub genes (20 hub genes each) from both the diseases, cluster 1 of RA and cluster C of CAD (Figure 5A). The STITCH-based interactions were derived from text mining, experiments, databases, co-expression, neighbourhood, gene fusion, co-occurrence, and prediction with a medium confidence level (0.400). To further validate our result in whole blood tissue and the extent of the applicability in RNA-seq, we specified the interaction to be constructed from the whole blood RNA-seq data retrieved from the Expression Atlas using the URL, https://www.ebi.ac.uk/gxa/baseline/experiments (Petryszak et al., 2014) with medium confidence level (0.400). It suggested a prominent connected component, GZMB, and some discrete nodes such as SASH3, and SLAMF8 (Figure 5A).

**Figure 5:**
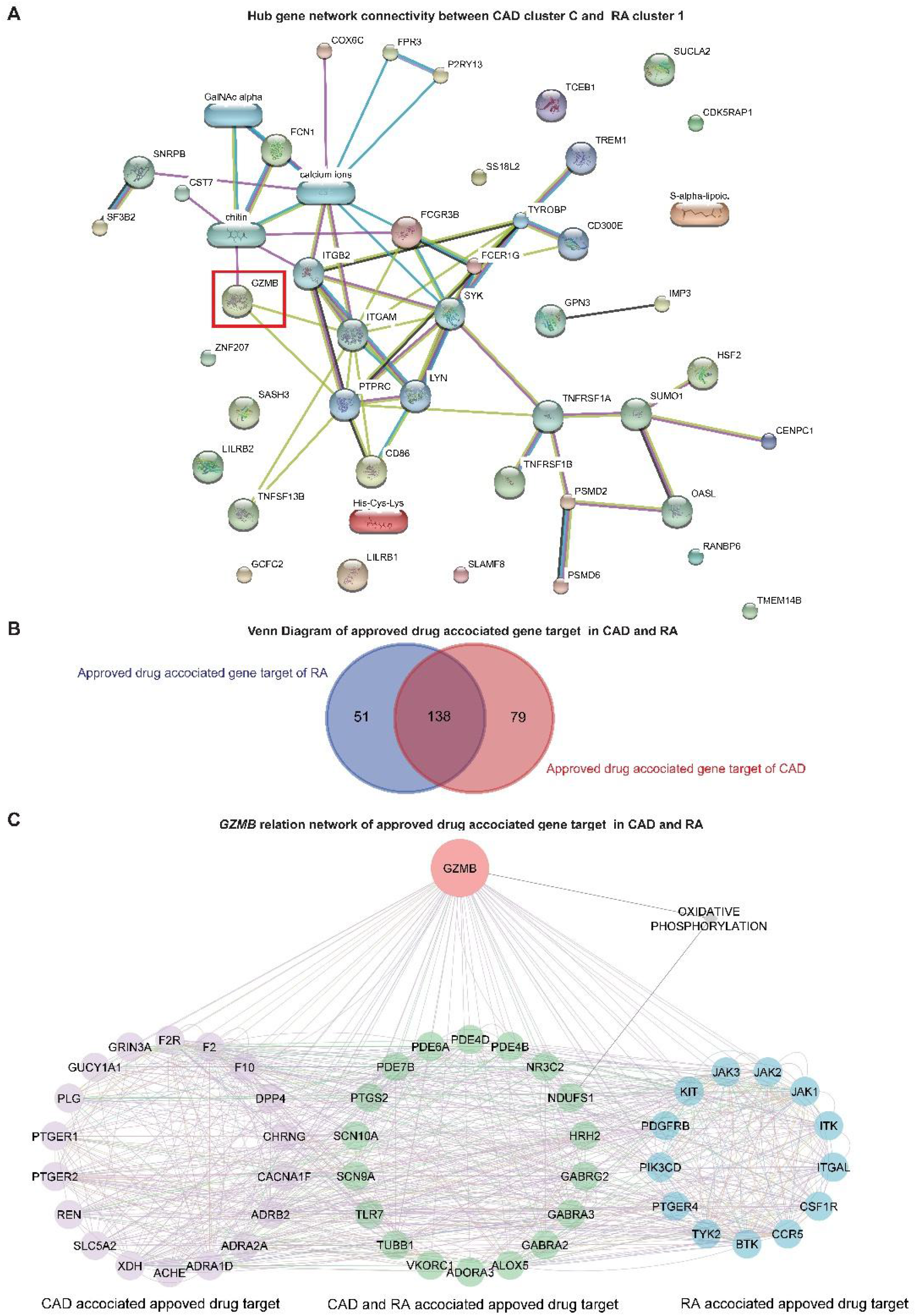
Network analysis and *GZMB*-related drugs targets. (A) The network analysis for the hub genes from cluster C of GSE59867 (CAD) and cluster 1 of GSE93272 (RA) hub genes using the STITCH database. The large nodes represent the known protein structures wherein the small nodes represent the unknown protein structures. The identified subpopulation-differentiation marker, GZMB was marked within a red box. The colours of the nodes and edges were predefined by STITCH. (B) The Venn diagram for the extracted drug-target genes (DTGs) of CAD (red) and RA (blue) using the Open Target platform. (C) Network-level association of *GZMB* and the related DTGs. The gene, *GZMB*, the ReDisX identified gene of interest (red), the CAD associated approved drug targets (purple), the approved drug targets for CAD and RA (green), and the approved drug targets for RA (blue) share a direct correlation. The colours of the edges were predefined by GeneMANIA.

### *GZMB*-related drugs target gene networks across CAD and RA

To evaluate the association of *GZMB* and its potential association with the drug target genes across CAD and RA, we employed the Open Targets platform (Koscielny et al., 2017). It facilitated the discovery of the available approved drugs and their association with the target genes (DTGs) of the diseases of interest, such as CAD and RA. A total number of 189 DTGs for RA and 217 DTGs for CAD were identified (Figure 5B). The Venn diagram analysis indicated that 73.02% of the RA-related DTGs intersected with CAD-related DTGs (Figure 5B). On the other hand, 63.59% of the CAD-related DTGs intersected with RA-related DTGs (Figure 5B). The detailed gene list is available in supplementary 4.

To investigate the network-level association of *GZMB* and those 268 unique DTGs (as shown in Figure 5B), GeneMANIA (Warde-Farley et al., 2010) was employed. The considered interactions were consolidated pathways, wiki-pathway, reactome, co-expression, physical interaction, drug-interaction, predicted, co-localization, pathway, shared protein domains and genetic interaction. This analysis extracted a total number of 36 additional genes out of the GeneMANIA knowledge base. Altogether it returned a total number of 305 genes and 18,138 interactions within the reconstructed network (supplementary 5). Further, we have narrowed our analyses by focusing on the *GZMB*-related DTGs in CAD and RA. It produced a network with the selected DTGs sharing direct interaction with *GZMB* (Figure 5C). This constructed network contains a total number of 51 genes and 755 interactions (Figure 5C).

## DISCUSSIONS

Diagnosis of a disease is used to get a universal consideration across the patients. Identifying the clinicopathological variables associated with the patients looks straightforward, but it is one of the trickiest and most sensitive concerns to be precisely addressed in terms of physiological and clinical perspectives. The current practices of classifying clinicopathological variables among the patients also referred to as classifications of diseases are typically relied on medical investigations such as physical and visual examinations, estimation of biochemical parameters, medical imaging methods, and of late a few genomic-markers-based diagnostic approaches, which are although not widely and commonly practised. Often the genomic disparity among the individual gets overlooked. It is something like a one-size-fits-all concept. However, with the advancement of the data-driven healthcare revolution and leveraging the high-throughput omics data, we could further investigate the issue as mentioned earlier to have better resolution in assessing the clinical conditions rather than a “typically classified disease and its same related treatments for all” for each individual. It may offer us a clue to identify the real root cause at the molecular layer and personalized or customized treatment plans. Our study supported our core rationale by introducing a functional concept of redefining diseases (with the working example of RA and CAD) based on individualized molecular characteristics using the ReDisX framework. Interestingly, our results indicated some exclusive subcategories featuring distinct molecular-genetic signatures within the “conventionally classified” disease category. ReDisX has tried to identify those distinct subpopulations upon thoroughly analyzing those signatures to reassign their clinical category focusing on precise identification.

### ReDisX offers better precision and personalized diagnosis strategy

Diagnosis of diseases usually relies on biochemical and pathological assessment, imaging techniques, and molecular analyses (Weigel & Dowsett, 2010). In many instances, especially for complex diseases, it is inadequate to explain the heterogeneity among the patients within a same disease and homogeneity across the patients from different diseases (Humby et al., 2019; Khera & Kathiresan, 2017; Niu et al., 2014). Therefore, redefining the diseases to elucidate the heterogeneity within a disease and homogeneity across the diseases will significantly ensure precision diagnosis and personalized treatments (Bell, 2010; Bezzina et al., 2022). Leveraging the advancement in acquiring high-throughput multi-omics data acts as an excellent indicator for the comprehensive molecular status of an individual, reflecting their health status and any impairments that occurred (T. Wang et al., 2021). So, it ensures precision and a personalized model of identifying patients’ clinicopathological conditions. Many computational and experimental studies have been evolved in this avenue (Aure et al., 2017; He et al., 2021; Li et al., 2015). They mostly attempted to address the heterogeneity within the patients under the same disease and homogeneity across the patients from different diseases. However, an interesting question remains unexplored about the extent of homogeneity amidst the heterogenous sub-population of different diseases. The consequent similarities and dissimilarities at the molecular level have not gained any comprehensive attention yet. Therefore, in this study, we have introduced ReDisX, a robust, scalable, and pathologically relevant computational framework to characterize the patients based on specific molecular-genetic signatures. We have systematically deployed the ReDisX framework considering two disease cases, RA and CAD. We have analyzed their transcriptional profiles to characterize the pathological similarity of their subpopulation. In the future, it perhaps guides us to extend this foundation to apply to other diseases to redefine their clinicopathological status aiming toward precision diagnosis and identifying personalized targets for therapeutic interventions.

Our proposed framework, ReDisX could differentiate the disease heterogeneity among the patients by optimally clustering them based on individual gene expression profile. One of our past studies indicated that RA and CAD used to share inflammatory pathways (Niu et al., 2014), and both of them consist of several subtypes (Khera & Kathiresan, 2017; van Wietmarschen et al., 2012). However, the detailed characteristics of the subpopulations of both RA and CAD were not yet fully explored. Our data suggested that the ReDisX could identify a distinct subpopulation in RA and CAD that were susceptible to mispronounced by the standard diagnosis criteria (as the data was captured originally at the point of diagnosis) without the ReDisX-based recategorization (Figure 2A, 2B, 3A, 3B). On the other hand, our results also indicated an identified homogeneity across the subpopulation of CAD to the RA patients (Figure 4, 5A). These characterizations were also validated using two other validation datasets strengthening our hypotheses (Figure 4A, supplementary 3). The aforesaid heterogeneity and homogeneity were not identified with the original diagnosis strategy mentioned in the dataset features. Therefore, the conventional modes of diagnosing diseases and categorizing the patients are used to overlook certain minute discrepancies at the individual molecular-genetic signature level. It may indicate a potential cause for failure of diagnosis of those specific subpopulations of patients, and therefore their receiving treatments might induce either inefficiency or some adverse effects. In many acute or severe clinical conditions, such inadequacy in characterizing the proper root cause might significantly delay the treatment process or prognostic outcomes. On this verge, our proposed framework, ReDisX showed an enhanced efficiency in accessing minute details of the molecular signatures in characterizing the patients.

### ReDisX supports advancing the screening of precise druggable target genes

Based on the ReDisX framework, our results suggested that *GZMB* was under-expressed in the subpopulation of both the diseases RA and CAD (Figure 4). A study by Joehanes, R. et al. concluded that the CAD patients with *GZMB* expressions exhibited a significant negative fold change (FDR<0.05) compared to their healthy controls (Joehanes et al., 2013). Although there was no solid evidence showing the direct pathological relationship between the *GZMB* under expression and the occurrence of CAD and RA, we have identified some diseases linked to *GZMB and* correlated with CAD and RA. For example, CAD was found to be correlated with Moyamoya Disease (MDD) (Peng et al., 2019), and pneumonia (Garcia-Laorden et al., 2016). Furthermore, Xing Peng et al. discovered that MDD patients with downregulated *GZMB* were also enriched with the downregulated genes of CAD (Peng et al., 2019). Many other studies also supported a correlation between MDD and CAD (Livesay & Johnson, 2019; Nam et al., 2015). Concerning the RA cases, a recent report showed that a patient with a history of RA for 15 years also suffered from cerebral rheumatoid vasculitis (El-Sudany et al., 2021). In a meta-analysis of 23 studies, it was found that the patients with RA shared a higher risk (∼1.68 times) of hemorrhagic stroke than normal individuals (Wiseman et al., 2016). This suggested that *GMZB-* may be a risk factor for RA patients developing MDD, but further investigation should be conducted to support this hypothesis. It also intrigued a sense of investigating *GZMB* as a potential drug target, especially for the niche subpopulation identified across the RA (cluster 1) and CAD (cluster C) patients wherein *GZMB* was under-expressed.

Studies also showed that excessive *GZMB* is related to inflammation (Francesca Velotti et al., 2020), a common pathological characteristic of CAD and RA (Golia et al., 2014; Shen et al., 2016; Sweeney & Firestein, 2004). However, those investigations did not consider minute fluctuations observed within some subpopulations of the CAD and RA patients who significantly exhibited *GZMB* under expression. Interestingly, our results characterized those distinct subpopulations of both CAD and RA as having *GZMB* under expression (Figure 4H, 4I). This claim was also validated with another RA dataset in our study (Figure 4J). This indication was also supported by the study conducted by Joehanes, R. et al. (Joehanes et al., 2013). The inhibition strategy for *GZMB* overexpression cases was considered a new therapeutic target for CAD and RA. Yue Shen et al. demonstrated that the under expression of *GZMB* protected against Ang II-induced cardiac hypertrophy and cardiac fibrosis, microhemorrhage, inflammation, and fibroblast accumulation (Shen et al., 2016). Cui-Xia Bao et al. show that *GZMB* gene silencing inhibits the MAPK signalling pathway by regulating the expressions of inflammatory factors (García-Laorden et al., 2016). Inherently, this demands our attention to investigate further those subpopulations of *GZMB* overexpression and underexpression and associated different clinicopathological statuses. After distinguishing its expression profiles within the subpopulation, it also sparked an idea to differentially employ *GZMB* as a drug target for personalized therapeutic intervention. Altogether, it essentially re-emphasizes that it is necessary to redefine the disease to navigate the hidden heterogeneity and cross disease homogeneity to offer better precise categorization, diagnosis, and treatment for the patients.

### ReDisX endorses the *GZMB* as a robust prognostic subpopulation differentiation marker to devise a personalized druggable target

GZMB (granzyme B) is a serine protease encoded by the *GZMB* and is commonly found in the natural killer (NK) cells and cytotoxic T cells (CTLs) (D’Eliseo et al., 2010). It has been reported to take part in different inflammatory signalling pathways such as inducing cell death, apoptosis (Choy, 2010), and suppression of viral replication (Afonina et al., 2010). It has also been reported as a drug-gene target for tumour therapy (Kurschus & Jenne, 2010), especially with cisplatin (Wishart et al., 2006) and mannose (Wishart et al., 2006).

However, there is no clinically recognized universal baseline for the *GZMB* expression pattern within the same or different diseases. Overexpression of *GZMB* was found to promote CAD (Ikemoto et al., 2009; Santos-Zas et al., 2021), wherein under-expressed *GZMB* was correlated with CAD (Joehanes et al., 2013) pathophysiology. It is a contradictory role of *GZMB* in CAD. Similar contradictory functions of *GZMB* were also reported in RA patients. For instance, the up-regulated *GZMB* was reported as an indicator of inflammatory diseases such as RA (F. Velotti et al., 2020), and in another research, the downregulation of *GZMB* by shRNA-mediated silencing was shown to promote RA in a rat model (C. X. Bao et al., 2018). It was found that *GZMB* silencing inhibited the MAPK signalling pathway by interfering with the expression of several inflammatory factors such as bcl-2, caspase and certain angiogenic factors such as VEGF, and bFGF. Indeed, a gene and its contradictory behaviours instigate an ambiguity in prognosis and consider that candidate for therapeutic development. It induces a layer of obscurity about devising inhibitors or activators to modulate *GZMB* within the same diseases and for different diseases, such as RA and CAD. Such complications represent the lack of understanding of the heterogeneity of *GZMB* across the subpopulations. Another recent single-cell study has also shared a similar view to consider the clinical identifications of RA based on different subsets of markers and reported the potential of *GZMB* as a subpopulation differentiation marker (X. Wu et al., 2021). So, it implies a solid motivation to utilize this gene, *GZMB*, as a prognostic marker or therapeutic target on a personalized basis. Added to their findings, determining the allocation of those markers across the RA patients at a personalized level would undoubtedly be an essential direction. The extent of the variations and discrepancies in their expression patterns prevails across the distinct subpopulation of RA patients is a prominent study scope. It will be even more interesting to navigate such similarities across the subpopulation of different diseases or dissimilarities within the same disease. Interestingly, our proposed framework, ReDisX addresses the expression pattern variation of *GZMB* across the different subpopulations of patients.

In our study, the ReDisX framework validates the potential of *GZMB* as a subpopulation differentiation marker within the same disease, e.g., under expression of *GZMB* in cluster 1 of RA patients and over expression of *GZMB* in clusters 2, and 3 of RA patients. Furthermore, it also demonstrated the homogeneity of *GZMB* expression across different diseases. For example, the under-expression pattern of *GZMB* of cluster 1 of RA patients shared a similar expression pattern with cluster C of the CAD patients, further supported by the functional enrichment analysis. These disparities strengthen the indication of the sensitivity of the strategies for therapeutically modulating *GZMB* expressions. Thus, it is highly recommended not to use a *GZMB* inhibitor/activator for all the patients suffering from RA and CAD, respectively.

A study reported the generic use of *GZMB* inhibitors to manage RA (C. X. Bao et al., 2018). However, our study flags a concern here and suggests the use of *GZMB* inhibitor to be devised on a personalized basis upon evaluating its specific patterns within the patient subpopulation. Similarly, it can also apply to CAD patients too. Additionally, the ReDisX framework suggests the specific drug designated for a subpopulation of RA patients with *GZMB* under-expression can be repurposed for the subpopulation of CAD patients having *GZMB* under-expression but never be used for the subpopulation of RA patients with *GZMB* overexpression. Hence, ReDisX demonstrated a data-driven ability to designate the *GZMB* as a potent subpopulation differentiation marker for RA and CAD. It also indicated a way to precisely deploy them for therapeutic development and a plausible strategy to repurpose those therapeutics backed by personalized gene-expression data.

### ReDisX suggests *GZMB* as a strategic focus for drug repurposing

*GZMB* inhibitors have been reported as therapeutics to manage inflammations related to RA (C. X. Bao et al., 2018) and CAD (Saito et al., 2011; Zeglinski & Granville, 2020). However, the contradictory dual role of *GZMB* has made the discovery process and its clinical application ambiguous (H. Wang et al., 2021). Studies have reported that *GZMB* overexpression and underexpression are linked to different clinical conditions, such as RA, CAD, and MDD (C.-X. Bao et al., 2018; Peng et al., 2019; Saito et al., 2011). It certainly makes the strategic development of therapeutics and/or discovering drugs against *GZMB* challenging. On the other hand, *GZMB* has been repeatedly endorsed as a prominent prognostic marker for many inflammatory pathways, especially related to RA, CAD, and angiogenesis (C. X. Bao et al., 2018). Hence, having a straightforward strategy to deal with *GZMB* is essential. ReDisX-based indication on *GZMB* and its prominent role as a prognostic marker has been further strengthened by the Open Target analyses (Figure 5C). It intensifies that the ReDisX-based identification is not only computationally validated, instead, it has a solid connection to be a potential personalized druggable target. It has also gained scientific support from different experimental studies and a presumed knowledge base (see result section for details). It motivates us to devise a strategy to deal with those ambiguities. So, ReDisX perhaps offers a plausible solution to designate the discrepancies of *GZMB* to be deployed as a prognostic marker and a target for therapeutic development.

### ReDisX is the robust, scalable, reproducible framework

Our study introduced ReDisX to characterize the heterogeneous subpopulations of a disease efficiently. It further considers additional molecular information about the gene expression, functional enrichment, druggable target genes, and homogeneity accessing layer. This has elucidated the deep expressional and pathological similarity of the subpopulations between two different diseases. We expanded our computational framework to reconstruct the hub gene network and identified their potential drug relevance. ReDisX indicated the heterogeneous subpopulations and the interconnected network of their DEGs. It also helped to quantify the homogeneity across different diseases, indicating two or more diverse clinical conditions sharing the similar underlying molecular dysfunctionality.

In this study, RA and CAD were used as an example stretching in connection with one of our prior studies (Niu et al., 2014). We have established our ReDisX framework as a proof-of-concept. It is scalable and can be deployed to any other disease context providing the input data are in the same format (as described in section 2). Methodologically, the infrastructure of ReDisX considers the ward’s distance in Hierarchical clustering and graph connectivity of the CMF model, wherein the ward’s distance is an established formula, and the hyperparameters in the CMF model were chosen as stated in the original paper (Wei et al., 2017; Yin & Tai, 2018). Hence, the reproducibility of the ReDisX could be ensured as long as the pre-labelling seeds are consistent.

The current practices of molecular pathological studies analyze the tissue-of-origin for any diseases (Hoadley et al., 2018). For example, the diagnosis of RA is typically made from synovial tissues (Fueldner et al., 2012) wherein cancers are diagnosed from tissue biopsies in the case of solid tumours (Wasserstrom et al., 1982). However, liquid biopsies are being widely practised (Russano et al., 2020). Eventually, the sampling procedures within clinical setup are gradually inclining towards non-invasive or minimally invasive ways while capturing the maximum information of the concerned underlying pathological conditions (Russano et al., 2020). So, keeping that vision, we put our effort to establish our model with the blood sample which is comparatively less invasive (Maciejak et al., 2015; Tasaki et al., 2018; Teixeira et al., 2009) and easier to be retrieved from patients and holds the potential to be a more expansive repertoire of the clinicopathological as well as molecular genetic conditions or biomarkers. However, based on our study objectives, we need to choose a suitable tissue to analyze all diseases. There are a few potential issues such as whole blood (Anaparti et al., 2017), saliva (Tar et al., 2021), and the gut microbiome (Shreiner et al., 2015) those are suitable for our study. In the future, the ReDisX framework could be extended and improved using multimodal data besides mRNA expressions such as methylation, and miRNA interference.

## CONCLUSION

ReDisX demonstrates a scalable data-driven framework to uniquely characterize the genomic signature and redefines the disease diagnosis strategy. It indicates a high-resolution precision and personalized diagnosis. It logically distinguishes the subpopulation heterogeneity within a disease and homogeneity across different diseases. It supports the personalized screening of DTGs. Our study with RA and CAD explains its efficiency in characterizing a subpopulation differentiation marker, *GZMB* augmenting it to be a potential personalized druggable target. Discovering the RA-characteristics-dominant CAD subpopulation supports one of our primary intentions to redefine the disease diagnosis with personalized molecular signature. It offers a new insight to understand the disease and revise our consecutive treatments plans. In addition, this study also suggests *GZMB* as a strategic focus for drug repurposing.

ReDisX framework is scalable and methodologically flexible to be further improved. We desire to deploy it to other complex disease cases and enhance the core algorithms by incorporating multimodal data. However, the clinical prospects of disease redefinition by ReDisX are yet to be validated in terms of the quality of precision diagnosis and efficacy of the recommended repurposed drug candidates against the indicated clinical conditions. Last but not least, it elucidated a novel perspective to rethink diagnosing diseases and the emergence of personalized therapeutic development.

## STRENGTHS OF OUR STUDY

1. ReDisX framework is a robust and scalable ML model applicable to any other disease of interest and can accommodate multimodal physiological high-throughput data.
2. It is a first-in-class CMF-based ML algorithm that precisely discovers signature gene expression features from the given patient data, and also, the small sample size does not affect the accuracy.
3. It discovers distinct heterogeneous subpopulations within a disease and homogenous subpopulations across different diseases.
4. It offers a clue to discovering new therapeutic targets and drug repurposing.
5. It can adopt a small sample size and return accurate predictions, consistent with many experimental studies.

## LIMITATION OF OUR STUDY

1. Hyperparameters used in ReDisX are data sensitive. It demands fine-tuning for different input data from different diseases.
2. Removal of cross-platform batch effect is susceptible to affect the accuracy of the analyses. Thus, it may require very careful preprocessing of input data.
3. The study’s current proposition, including mRNA gene expression data, could partially explain the body’s underlying mechanism.
4. The proposed framework of ReDisX is based on a single omics analysis algorithm, and its impact in the case of multi-omics input across multi-tissue data is not optimized yet.
5. The generalization of our proposed method is still subjected to be validated before any clinical applications.

## MATERIALS AND METHODS

### 1. Dataset and preprocessing

#### 1.1. Gene expression data

Two publicly available datasets, GSE93272, and GSE59867 were collected from the National Center for Biotechnology Information Gene Expression Omnibus (NCBI–GEO). The GSE93272 is a gene expression data obtained from a whole blood transcriptome (Tasaki et al., 2018)(Tasaki et al., 2018). It contains 275 samples from RA patients and healthy controls, performed with Affymetrix Human Genome U133 Plus 2.0 (GPL570). The whole blood transcriptome dataset was retrieved from https://www.ncbi.nlm.nih.gov/geo/query/acc.cgi?acc=GSE93272. The RA patients data within this dataset was categorized into three groups: (1) the RA patients without any receiving treatments, (2) the RA patients receiving treatments such as methotrexate, infliximab, and tocilizumab, and (3) the healthy controls(Tasaki et al., 2018). The dataset contains a total number of 20,356 genes’ expression values for each patient which was further processed and normalized using Robust Multi-Array Average (RMA).

The GSE59867 is also a whole blood transcriptome dataset (Maciejak et al., 2015)(Maciejak et al., 2015). It contains 436 samples from the CAD patients and healthy controls, performed with Affymetrix Human Gene 1.0 ST Array (GPL6244). The data was retrieved from https://www.ncbi.nlm.nih.gov/geo/query/acc.cgi?acc=GSE59867. The samples were collected from peripheral blood from patients (*n*=111) with ST-segment elevation myocardial infarction (STEMI) at four-time points (admission, discharge, one month after myocardial infarction (MI), and six months after MI). The control group consists of 46 individuals either with reported stable CAD or without any history of MI. The dataset contains a total number of 20,511 genes’ expression values for each patient which was further processed and normalized using RMA.

#### 1.2. Data preprocessing

Intersections of the genes (*m*=17,432) were extracted from both the datasets, GSE93272 and GSE59867. Top 5,000 High Variance Genes (HVGs) were filtered out for further analysis, and the voom (Law et al., 2014) (Law et al., 2014)was applied separately to estimate the mean-variance relationship between those two datasets.

### 2. Construction of the ReDisX framework and applications

ReDisX is an ML framework primarily relying on a CMF model (Wei et al., 2017; Yin & Tai, 2018). The construction of the ReDisX framework consists of 2 main parts, (1) hierarchical clustering for pre-labelling all the sample points, and (2) part of the pre-labelling samples would be used in CMF for final labelling for each sample (please see section 2.5 for details) (Figure 1B).

#### 2.1. Clustering the patients using ReDisX

After sorting out the HVGs, the ReDisX was applied to cluster the patients based on gene expression similarity patterns (see section 2.2 for notation, section 2.3 for hierarchical clustering, and section 2.4 for the detailed construction of CMF model) (Figure 1B). Then, each patient was labelled where the same label represents a higher similarity discovered in their expression profile.

#### 2.2. Notations

We used lower-case letters such as *x* to represent scalar or single random variables, bold lower-case letters such as ***x*** to represent a vector variable, bold capital letters such as ***X*** and letters in calligraphy such as *𝒳* to denote a matrix variable and a set, respectively. For example, in this paper, the data set was denoted as a calligraphic capital letter ***𝒳*** = {***x***_1_, ***x***_2_, …, ***x***_*n*_} and *X* = ***x***_1_, ***x***_2_, …, ***x***_*n*_) ∈ ℛ^*d*×*n*^ is a matrix of the aligned vectors, respectively, while ***x***_***i***_ are *d*-dimensional variables, and *n* is the number of the data samples, ***x***_*i*_ It is a data point of the *i*-th sample, denoted as a vector. We also used ‖ · ‖ to measure the length of the vector, and it used Euclidean or *L*_2_ norm by default in the following part if there is no other explanation. In this paper, we used *n* and *K* as the number of samples in the dataset and the number of clusters to be divided, respectively. The notation *𝒞* was used to denote the dataset and *𝒞*_*k*_ represents the *k-th* cluster. Normalization of the matrix ***X*** concerning dimension was performed as a prerequisite to executing the normalization function in MATLAB.

#### 2.3. Hierarchical clustering

Agglomerative hierarchical clustering is one of the most popular clustering methods. The detail was investigated by (Day & Edelsbrunner, 1984)(Day & Edelsbrunner, 1984). It aims to construct a hierarchical partition of data samples using a greedy strategy. In the initial stage, all data samples were viewed as individual cluster. Each turn takes a pair of two nearest clusters and merges them into a larger cluster. Thus, the number of clusters used to get decreased by one after each turn. MATLAB has also implemented agglomerative hierarchical clustering in some **clusterdata** and **linkage** functions. In our experiments, we used Ward’s method, which is also known as the average linkage or minimum variance method, to measure the distance between two clusters, which is defined as follows,

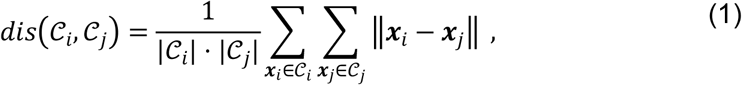

where |*𝒞*_*i*_| represents the number of samples in the *i*-th cluster.

#### 2.4. The Continuous Max-Flow (CMF) model

The graph models for clustering view the data points as individual vertix and use edge sets to describe the relation between different data points. This well-known model, CMF was constructed relying on the max-flow/min-cut theorem(Yin & Tai, 2018). It is used to depart the graph into *K-*connected components with minimal cutting edges. The continuous version of the max-flow/min-cut model was termed continuous max-flow or CMF which offered an efficient numerical method (Yin & Tai, 2018)(Yin & Tai, 2018). It minimized the cost of edge cutting and introduced a region force that could extract the preliminary information from the dataset with other techniques. We used the same specification as stated by (Yin & Tai, 2018)(Yin & Tai, 2018) in this study, and the region force term was defined in the followings,

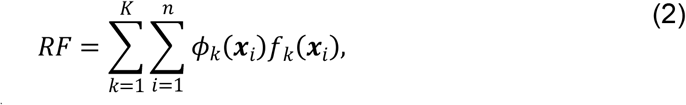

where 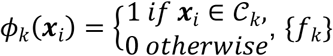, is the region forces. We set that *f*_*k*_(***x***_*i*_) = − *log (p*_*k*_ (***x*** _*i*_))+ *log* (1 − (*p*_*k*_ (***x*** _*i*_)) likes in (Yin & Tai, 2018)(Yin & Tai, 2018), where the function {*p*_*k*_} a set of prior conditional probabilities which has been discussed in section 2.5. The min-cut problem with region force was to minimize the following energy,

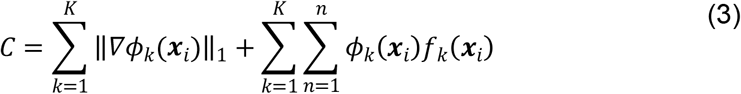

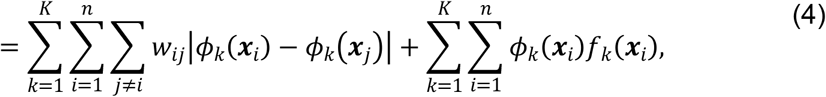

where 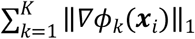 was defined as the term edge cutting, and {*w*_*ij*_} are the weights of the edge between the data points ***x***_*i*_ and ***x***_*j*_ which was discussed more in the following section 2.5.

Let (*ϕ*_1_, *ϕ*_2_, …, *ϕ*_*k*_) ∈ {0,1}^*n*×*K*^, and ***F*** = (*f*_1_, *f*_2_, …, *f*_*k*_) ∈ ℛ^*n*×*K*^, ***W*** =(*w*_*ij*_) ∈ ℛ^*n*×*K*^, *and* ***D*** = (***d***_*ii*_) ∈ ***R***^***n*** ×***n***^ would be explicitly defined in section 2.5, then the above cost function could also be written as the following,

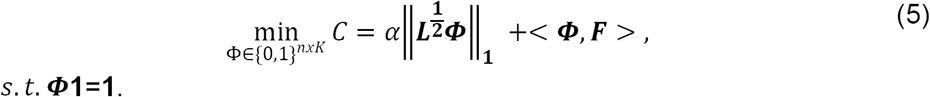

where ***L*** = ***W*** − ***D***, ‖***A***‖_1_ = Σ_*i,j*_| *A*_*ij*_ | and *α* is a regularization parameter that balances the region force and min-cut term. However, searching for the optimal solution ***Φ*** in discrete space {0,1}^*nxK*^ was an **NP**-hard problem, then the feasible space of the above problem was relaxed into a continuous one as eq. (5) and solved the following problem

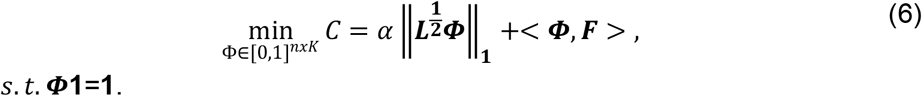

Then, this optimal problem becomes equivalent to the min-max problem as eq. (6),

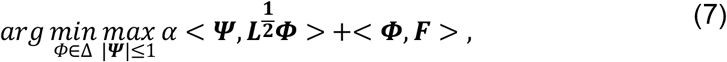

where Δ = {***Φ*** ∈ [0,1]^*n*×*K*^: ***Φ*1** = **1**}. We also used the primal-dual method with the projection technique to solve this problem. The details of the algorithm can be found in (Yin & Tai, 2018)(Yin & Tai, 2018).

#### 2.5. Weight function

We now describe how to evaluate a prior probability that each data point ***x***_*i*_ belong to the *k*-th cluster *𝒞*_*k*_, denoted as *p*_*k*_(***x***_*i*_) in section 2.4. In the pre-labelling stage, each datapoint *x*_*i*_ were classified into the *k*-th cluster by Hierarchical method. Among those data points, part of the would be used as a pre-label for the CMF model, which would be assigned *p*_*k*_(***x***_*i*_) = 1 and *p*_*j*≠*k*_(***x***_*i*_) = 0. The pre-label would be equal for each cluster, and the number of pre-labels for each cluster would be the minimal number of elements in all clusters. For other data points, we introduced a metric on the data that measures the rate of connectivity of the points ***x***_*i*_ and ***x***_*j*_ based on the m-steps diffusion process (Coifman et al., 2005)(Coifman et al., 2005). The diffusion process could be viewed as a Markov chain, and in our experiment, the transition matrix ***A*** ∈ ℛ^*n*×*n*^, ***A***_*ij*_ means the probability of the data point diffuses from ***x***_*i*_ to ***x***_*j*_. Let 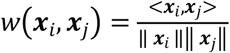, then we set 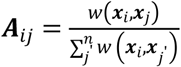. Denote ***D*** = (*d*_***ii***_) ∈ ℛ^*n*×*n*^is a diagonal matrix where 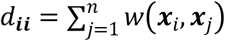, and 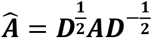 is a symmetric matrix. Then the m-step diffusion distance between *x*_*i*_ and *x*_*j*_ is defined as 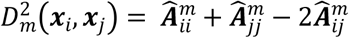.

Here 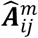 could interpret the transition probability through the m-step diffusion (while 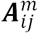 is the real probability), then we define the probability of ***x***_*i*_ belong to cluster *k* as

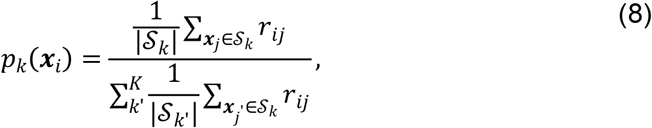

where *𝒮*_*k*_ is a set of labelled data samples in the *k*-th group which we discussed in the preprocessing part, |*𝒮*_*k*_| is the number of samples in the corresponding group, and 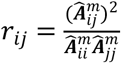 represents the similarity between ***x***_*i*_ and ***x***_*j*_. In an actual implementation, we chose *m*=2. Similarly, we also chose the cosine distance to measure the distance between ***x***_*i*_ and ***x***, and 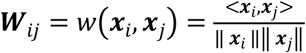. Considering the cost of computation, for each data point, we only use the edges between other *K*-nearest data points. We chose *K*=10 using the *K*-nearest neighbour algorithm (KNN), where *K* represents the cluster number.

#### 2.6. Cluster evaluation

As the core CMF model firmly considers the graph connectivity, it tends to merge two similar groups identified by Hierarchical clustering. Based on the merging cluster property, the ReDisX framework was deployed twenty times by iterating the number of clusters from 1 to 20. Further, the unique number of clusters was evaluated upon merging the graph connectivity as indicated by the CMF model. Then, the most occurrence number of clusters was chosen as the optimal number of clusters.

### 3. Screening the DEGs

The DEGs were extracted using the Limma package in R (Ritchie et al., 2015), considering the p-value <0.05 and log-fold change >0.05 for both the datasets GSE93272 and GSE59867. A subpopulation within both the datasets represents a set of DEGs about their respective healthy controls. Then, the DEGs were filtered out for both datasets to extract the overlapping genes across the other subpopulations. The sorted overlapping and non-overlapping genes were considered further for our analyses.

### 4. Functional enrichment analysis

The non-overlapping DEGs for both datasets were subjected to functional enrichment analysis. Those genes were further functionally characterized using Enrichr (Chen et al., 2013; Kuleshov et al., 2016) to discover the maximum enrichment for DisGeNet (Piñero et al., 2017) disease information and to have the most likely pathways using KEGG (Kanehisa & Goto, 2000) pathway analysis.

### 5. Knowledge-based DEGs network

In parallel to the functional enrichment analysis, a knowledge-based network extraction using GeneMANIA (Warde-Farley et al., 2010) was also performed for both datasets(Warde-Farley et al., 2010). So, a knowledge-based network consisting of the DEGs was constructed for each subpopulation within those two datasets.

### 6. Hub genes identification

After constructing and analyzing the knowledge-based networks, the hub genes within each subpopulation were identified. cytoHubba (Chin et al., 2014), a dedicated plugin under Cytoscape (v3.9.1) (Shannon et al., 2003), was used to discover the hub genes. In each subpopulation, 20 hub genes were identified.

### 7. Drug target screening

Those identified hub genes (from each subpopulation of both datasets) were examined against the drug bank deposited drug targets (Wishart et al., 2006). Upon screening the pool of hub genes to discover potential drug targets, they were subjected to fit the drug-related gene target matching.

### 8. Validation of the Hub genes

The ReDisX-based hub genes’ features, especially the co-expression patterns and expression patterns of the DEGs within those hub genes, were validated using two other publicly available datasets, GSE15573 (Teixeira et al., 2009) and GSE77298 (Broeren et al., 2015). The GSE15573 is a whole blood transcriptome dataset(Teixeira et al., 2009) containing 33 samples from RA patients and healthy controls obtained by Illumina human-6 v2.0 bead chip (GPL6102). The raw data was retrieved from https://www.ncbi.nlm.nih.gov/geo/query/acc.cgi?acc=GSE15573. Then, the GEO2R tool (Barrett et al., 2013) was applied to check the DEGs for validation. Similarly, another dataset, GSE77298, a synovial biopsies-derived transcriptomic dataset, (Broeren et al., 2015)consists of 16 samples from RA patients and healthy controls obtained using Affymetrix Human Genome U133 Plus 2.0 Array (GPL570). Raw data was retrieved from https://www.ncbi.nlm.nih.gov/geo/query/acc.cgi?acc=GSE77298. Then, GEO2R (Barrett et al., 2013) was also employed to check the DEGs for validation purposes.

#### Cross diseases subpopulation hub gene network analysis

STITCH (http://stitch.embl.de/) (Kuhn et al., 2014; Kuhn et al., 2008) database was applied to examine the hub gene network similarity discovered by cytoHubba (Chin et al., 2014). It also examines the interdependence among the hub genes across the subpopulation of different diseases. The interactions were identified through text mining, experiments, databases, co-expression, neighbourhood, gene fusion, co-occurrence, and prediction with a medium confidence level (0.400).

### 9. Dataset visualization

#### PCA & t-SNE

PCA is one of the most popular and efficient linear approaches for dimension reduction (Hotelling, 1933; Pearson, 1901)(Hotelling, 1933; Pearson, 1901). PCA’s singular value decomposition technique is an efficient method for extracting data features in low-dimensional linear subspace (Hotelling, 1933; Pearson, 1901). However, PCA exposed its limitations for high-dimensional data lying on or near a low-dimensional manifold. Hence, a non-linear technique called t-SNE (van der Maaten & Hinton, 2008) was also applied(van der Maaten & Hinton, 2008). As Van der Maaten suggested, t-SNE is also limited in high time complexity (van der Maaten & Hinton, 2008). In our study, the PCA and t-SNE were used in a combinational scheme to visualize the gene expression data from the patient population in a 2D way. MATLAB (v2021b) was used to execute the functions **pca** and **tsne** in this study.

### 10. Drug associated network analysis

Open Targets platform (https://www.opentargets.org/) (Koscielny et al., 2017) was employed for knowledge extraction for its association with the target genes across CAD and RA. Next, we narrowed down our target to the approved drugs only for the DTGs for rest of the analyses. Then, the network-level association of the ReDisX identified gene and those DTGs were analyzed using GeneMANIA (Warde-Farley et al., 2010). The considered interactions were consolidated pathways, wiki-pathway, reactome, co-expression, physical interaction, drug-interaction, predicted, co-localization, pathway, shared protein domains and genetic interactions.

## Author contribution

HFY and DC contributed to conceptualization, contextualizing, study designing, and refining. HFY and DC analyzed and interpreted the data, and also prepared the original draft and the figures of the manuscript. DC contributed to overseeing the project and revising the manuscript. XCT, KFL, and CZ contributed to constructing the algorithms and mathematical executions. KW, YL, YG, LL, and DG contributed to data interpretation. DG also supported the work. HLZ, XCT, and APL supervised and supported the work. All authors provided critical feedback and approved the submitted version.

## Data availability

## Acknowledgements

We sincerely thank the facilities at the School of Chinese Medicine, the Department of Mathematics of Hong Kong Baptist University, Hong Kong for providing the necessary support for this study.

