## Supplementary figures and images for "ReDisX: a Continuous Max Flow-based framework to redefine the diagnosis of diseases based on identified patterns of genomic signatures"

### Barplot_15573_59867_DisGeNet.pdf

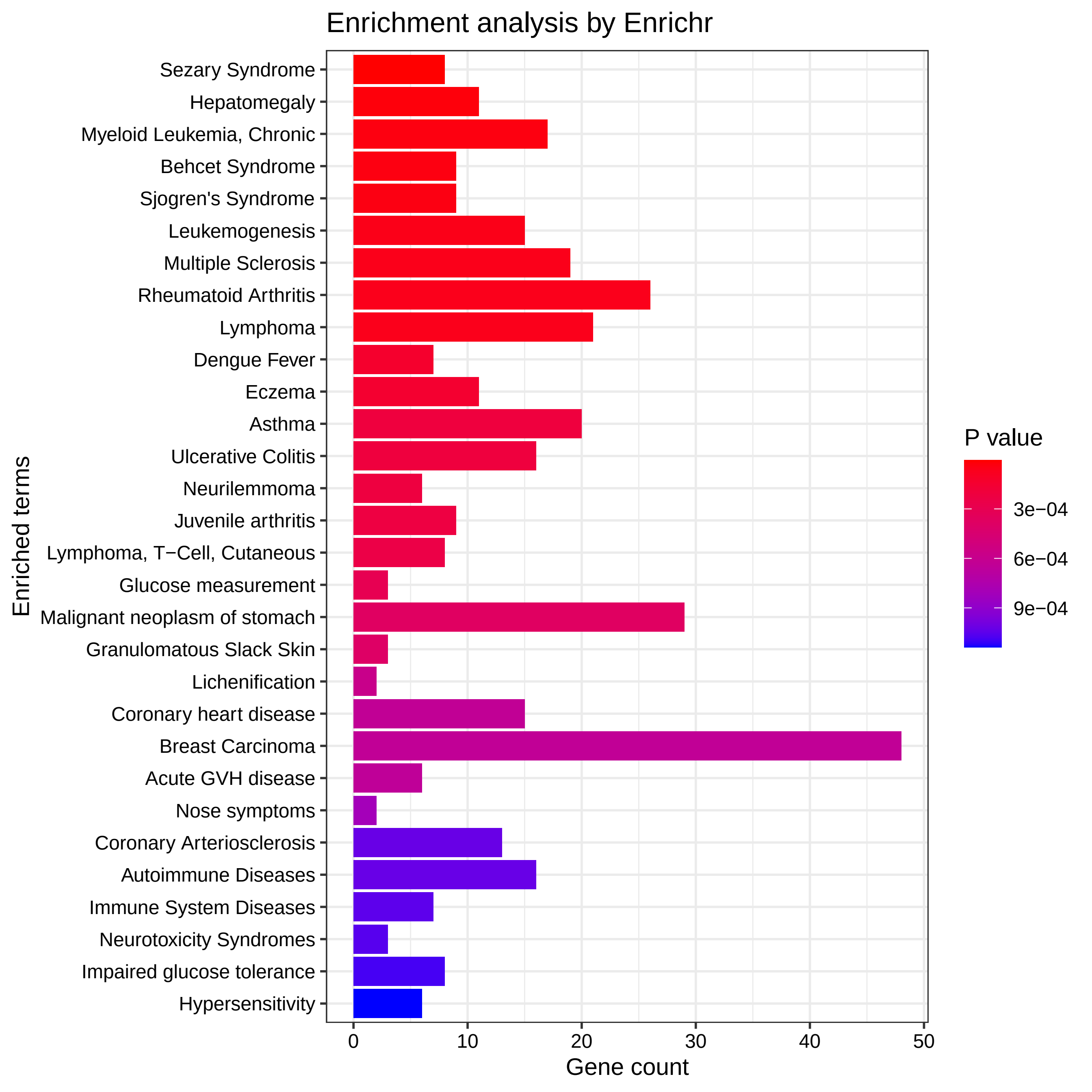

### Barplot_15573_59867_GO_BP.pdf

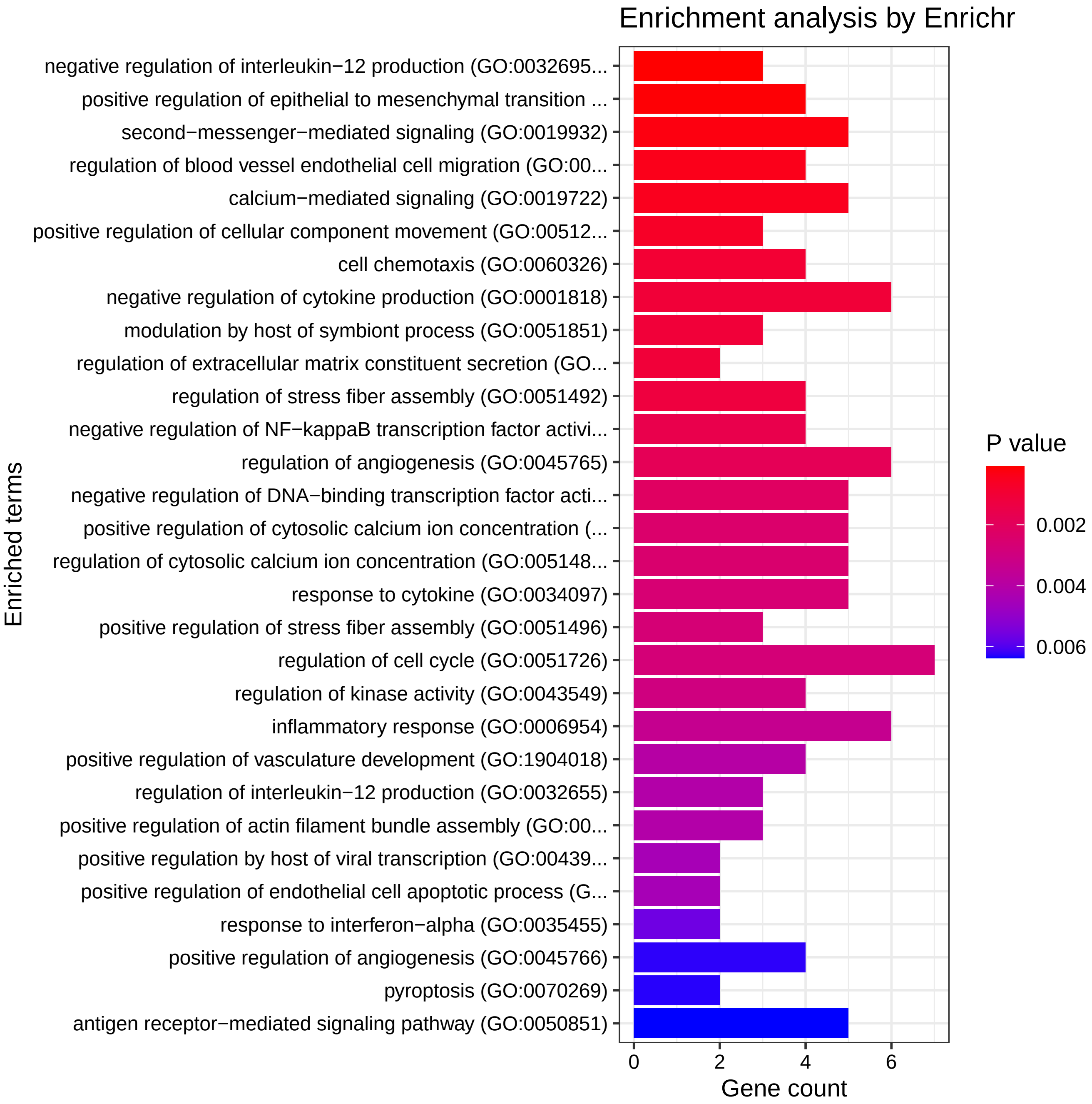

### Barplot_15573_59867_GO_CC.pdf

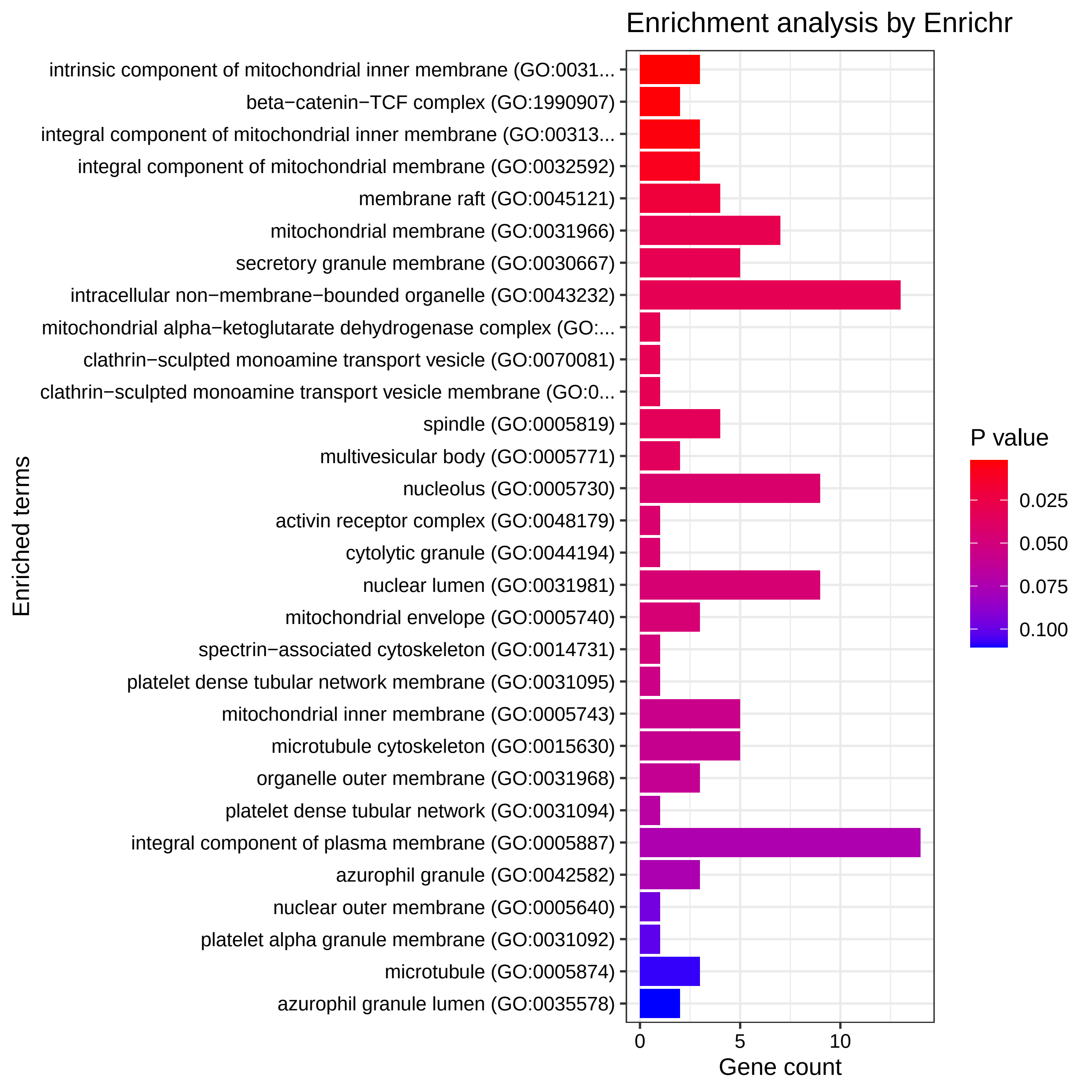

### Barplot_15573_59867_GO_MF.pdf

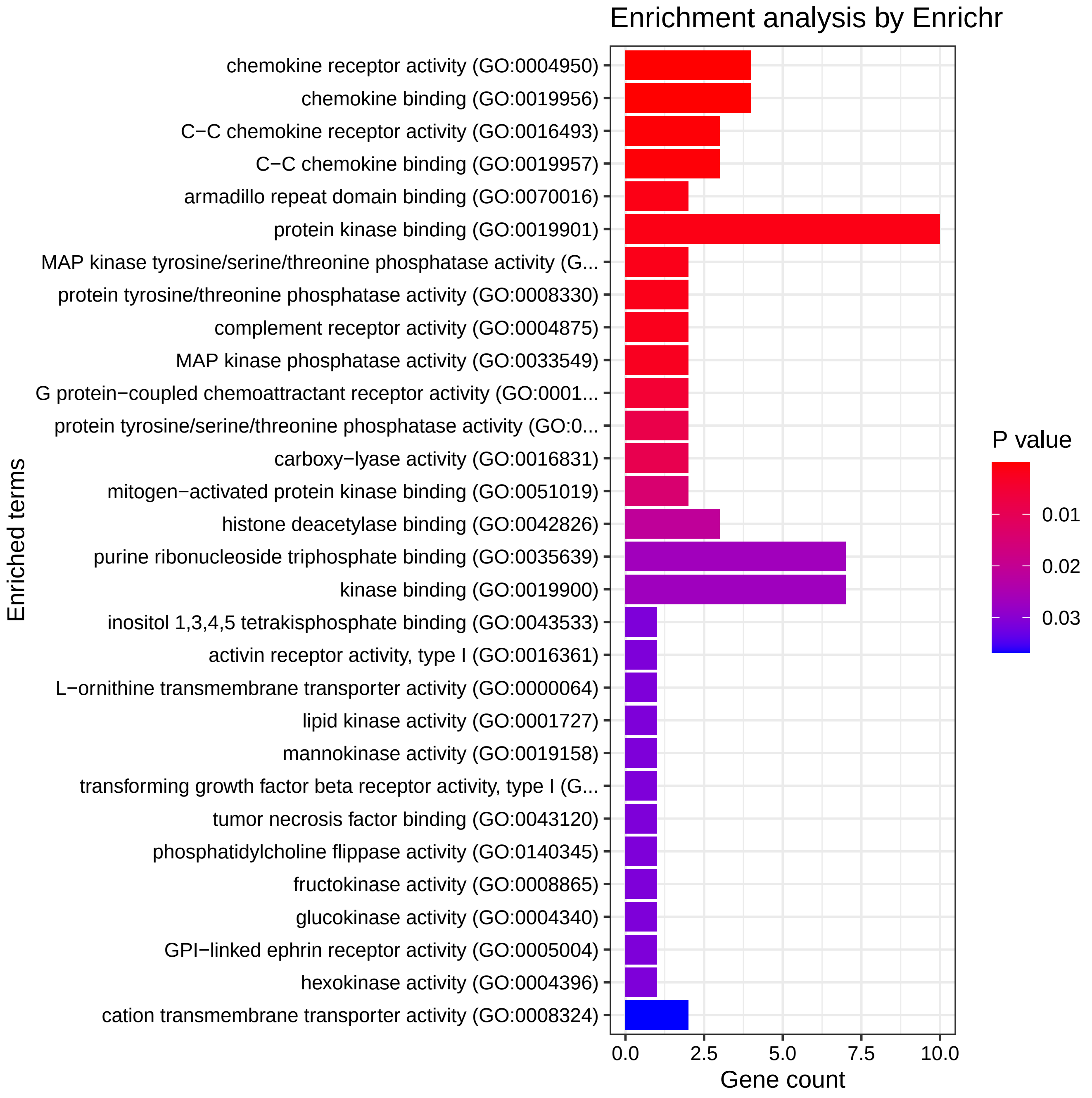

### Barplot_15573_59867_KEGG.pdf

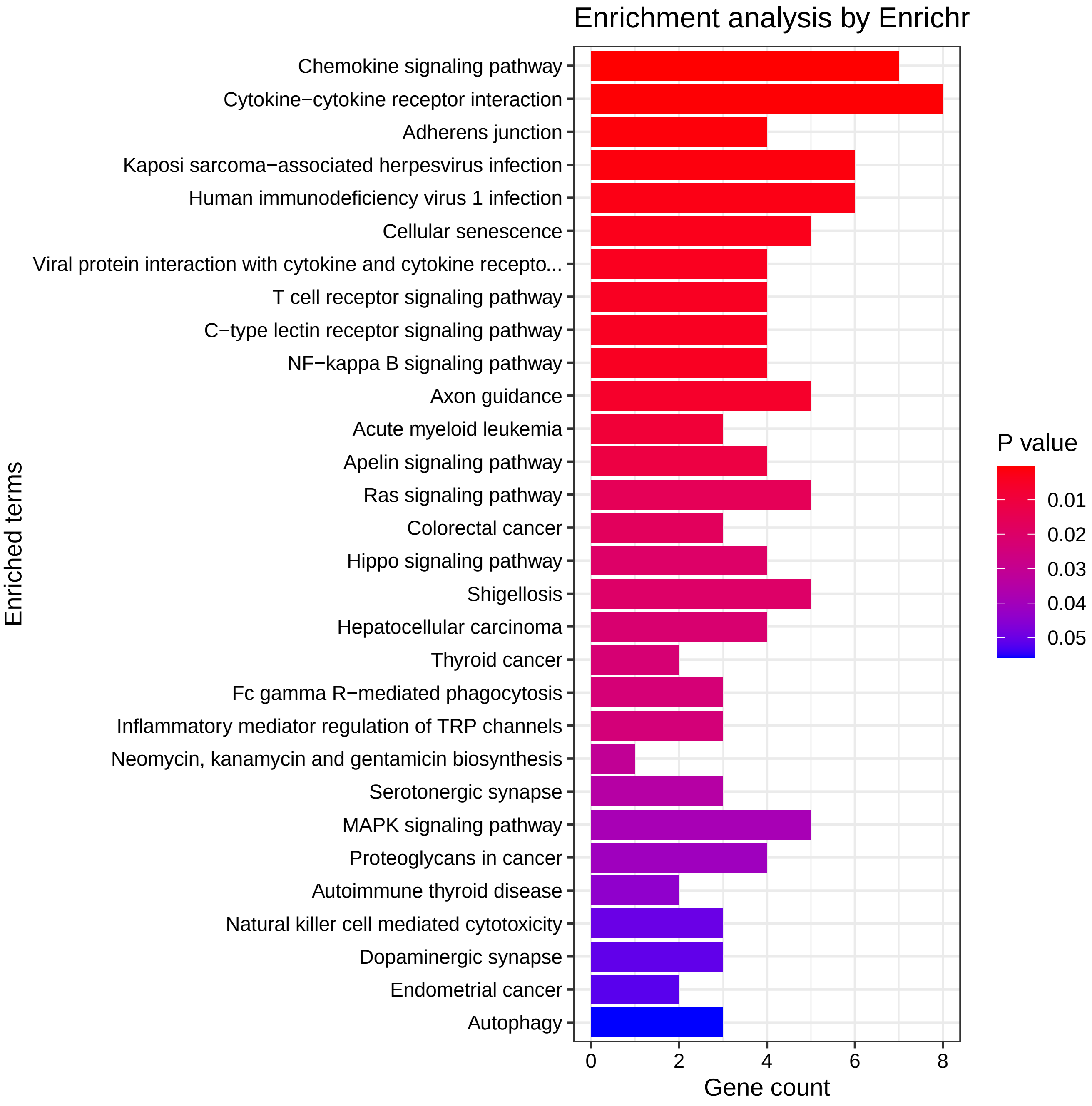

### Barplot_77298_59867_DisGeNet.pdf

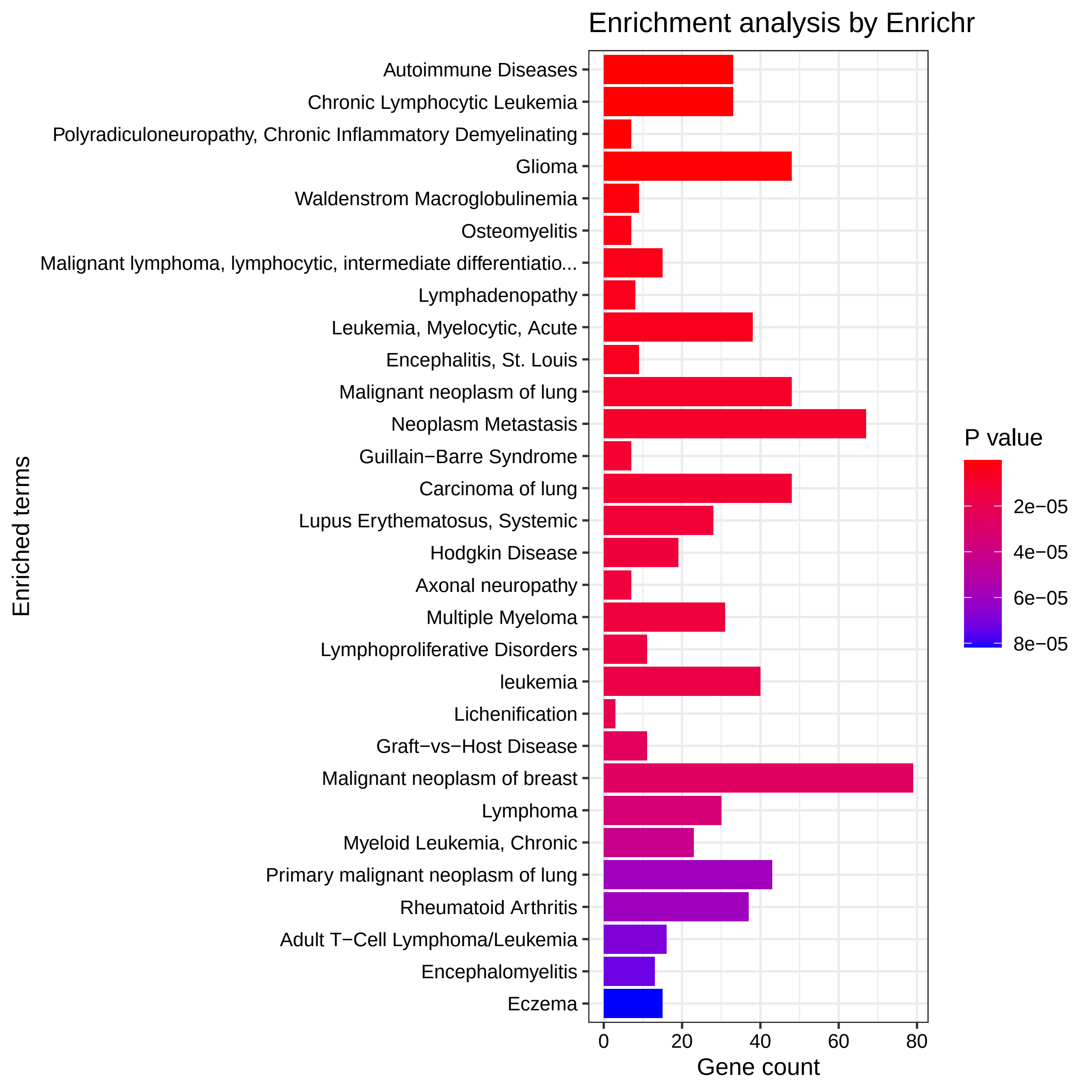

### Barplot_77298_59867_GO_BP.pdf

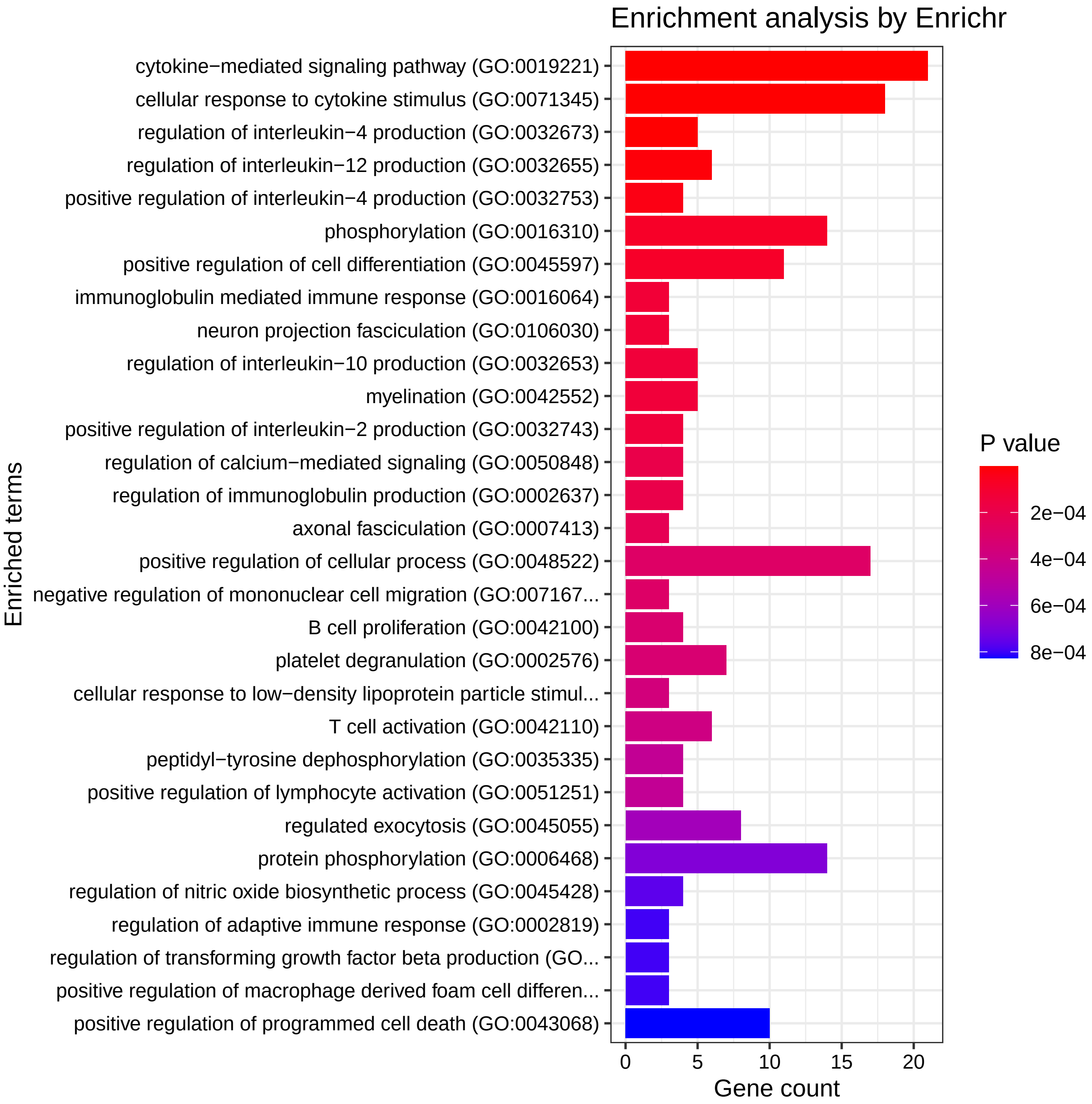

### Barplot_77298_59867_GO_CC.pdf

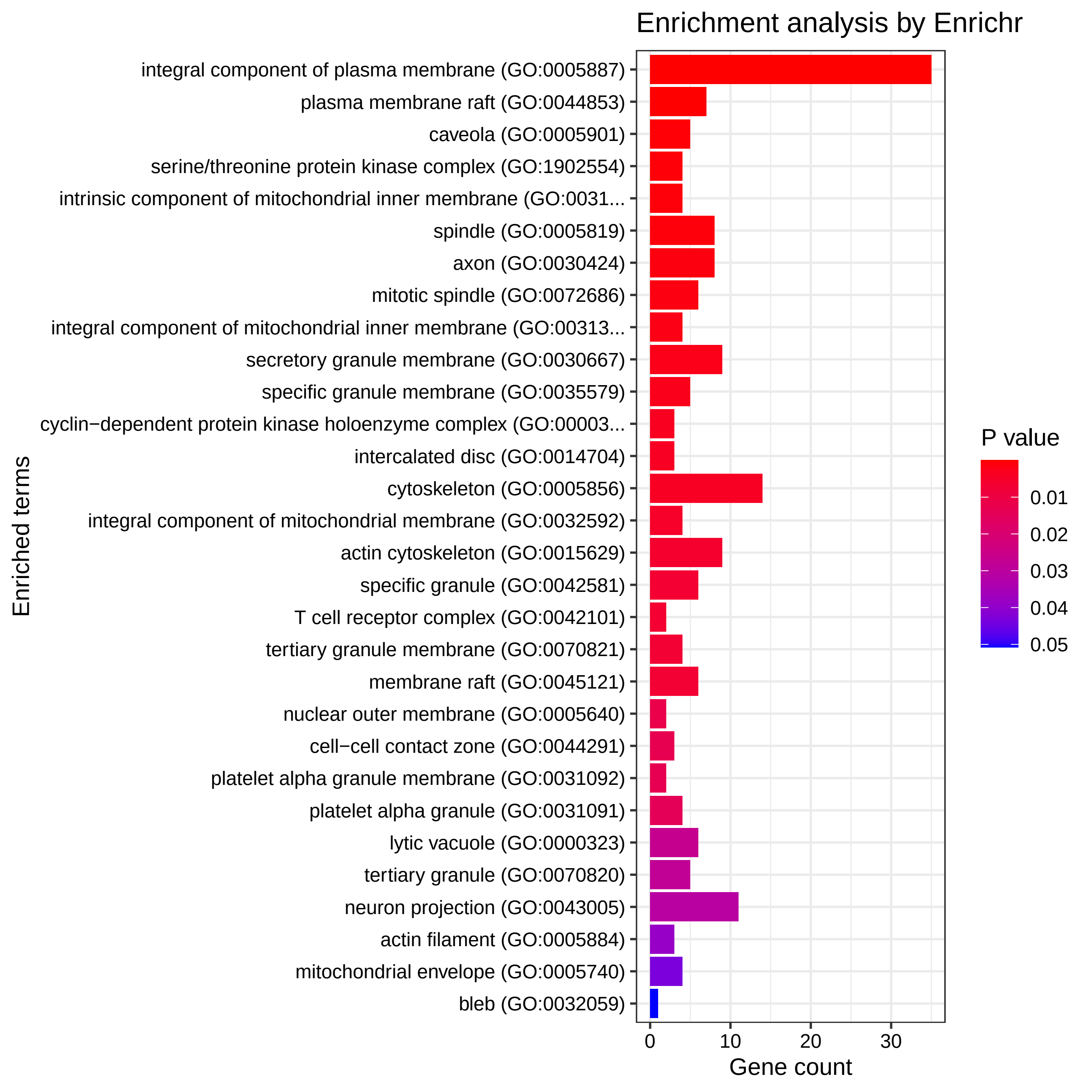

### Barplot_77298_59867_GO_MF.pdf

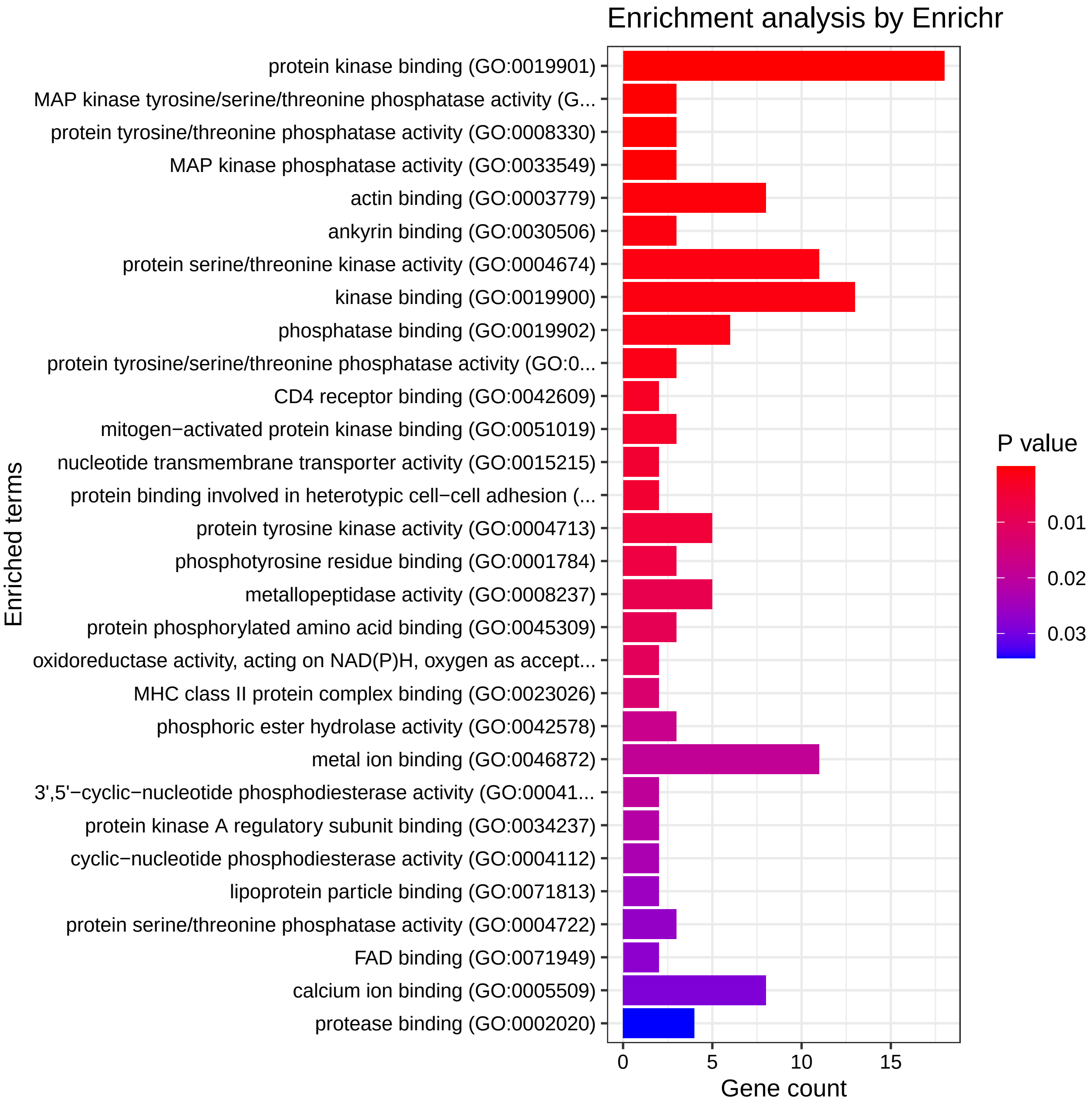

### Barplot_77298_59867_KEGG.pdf

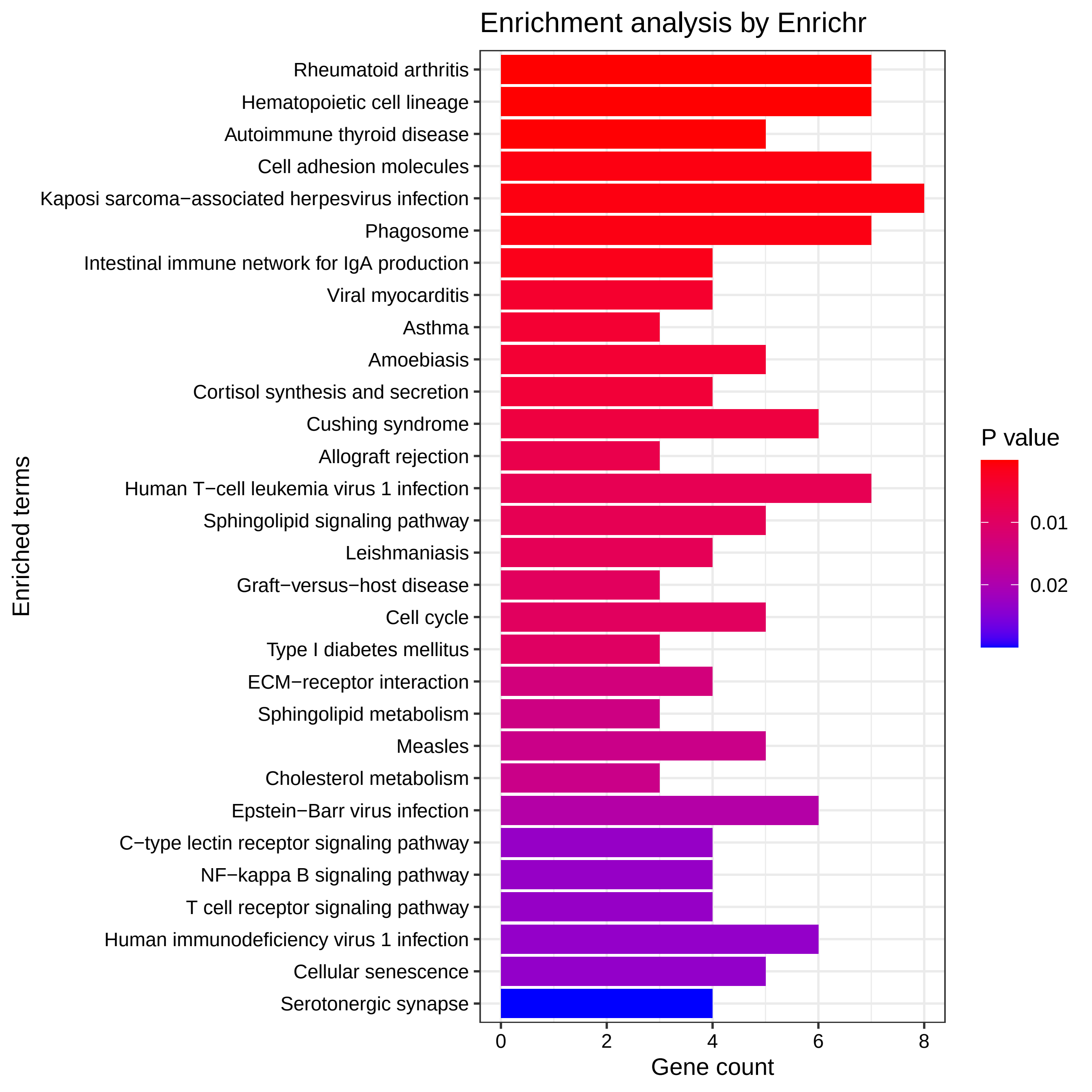

### Barplot_ReDisX_GSE59867_clus3_DisGeNet.pdf

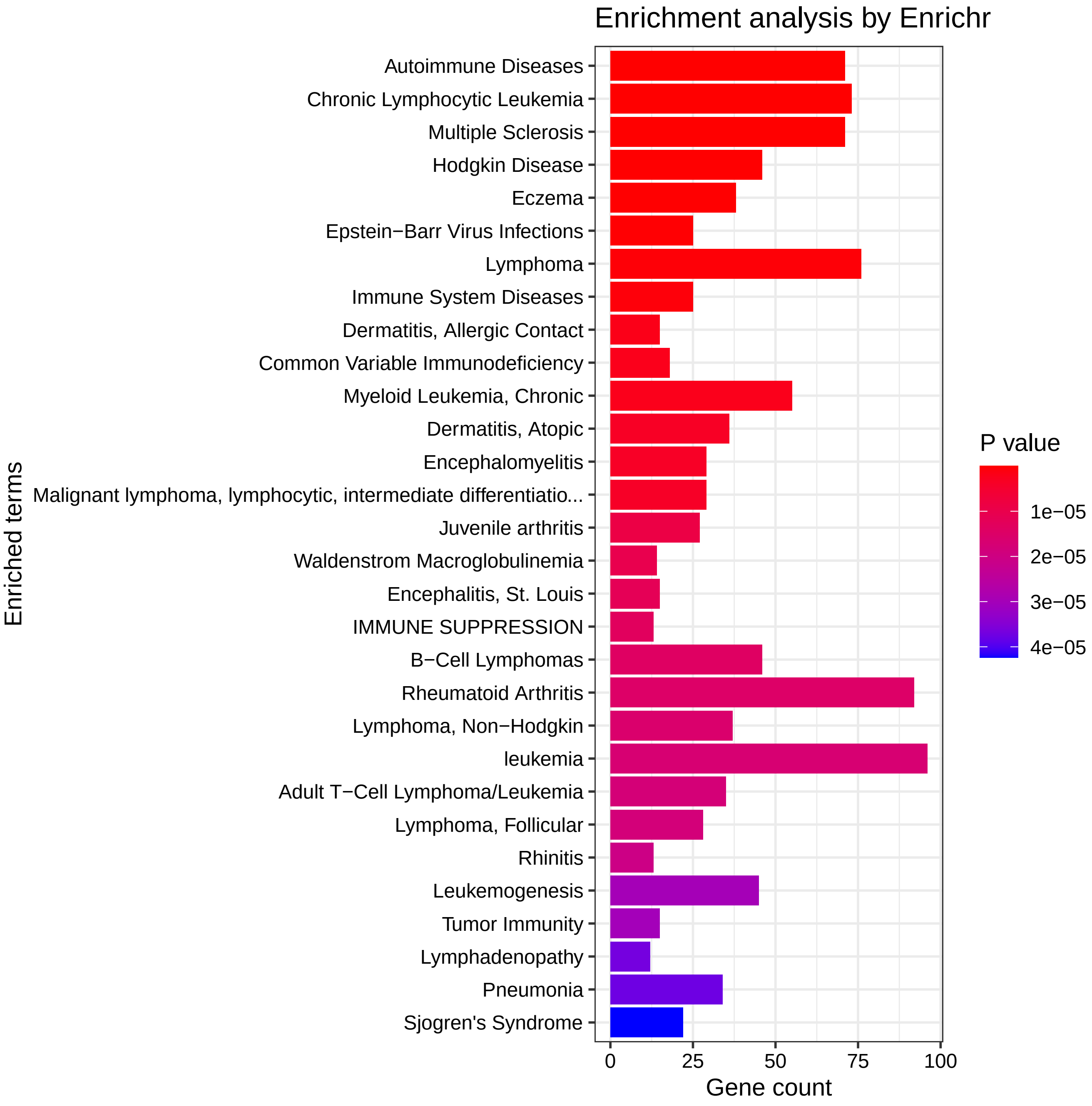

### Barplot_ReDisX_GSE59867_clus3_GO_BP.pdf

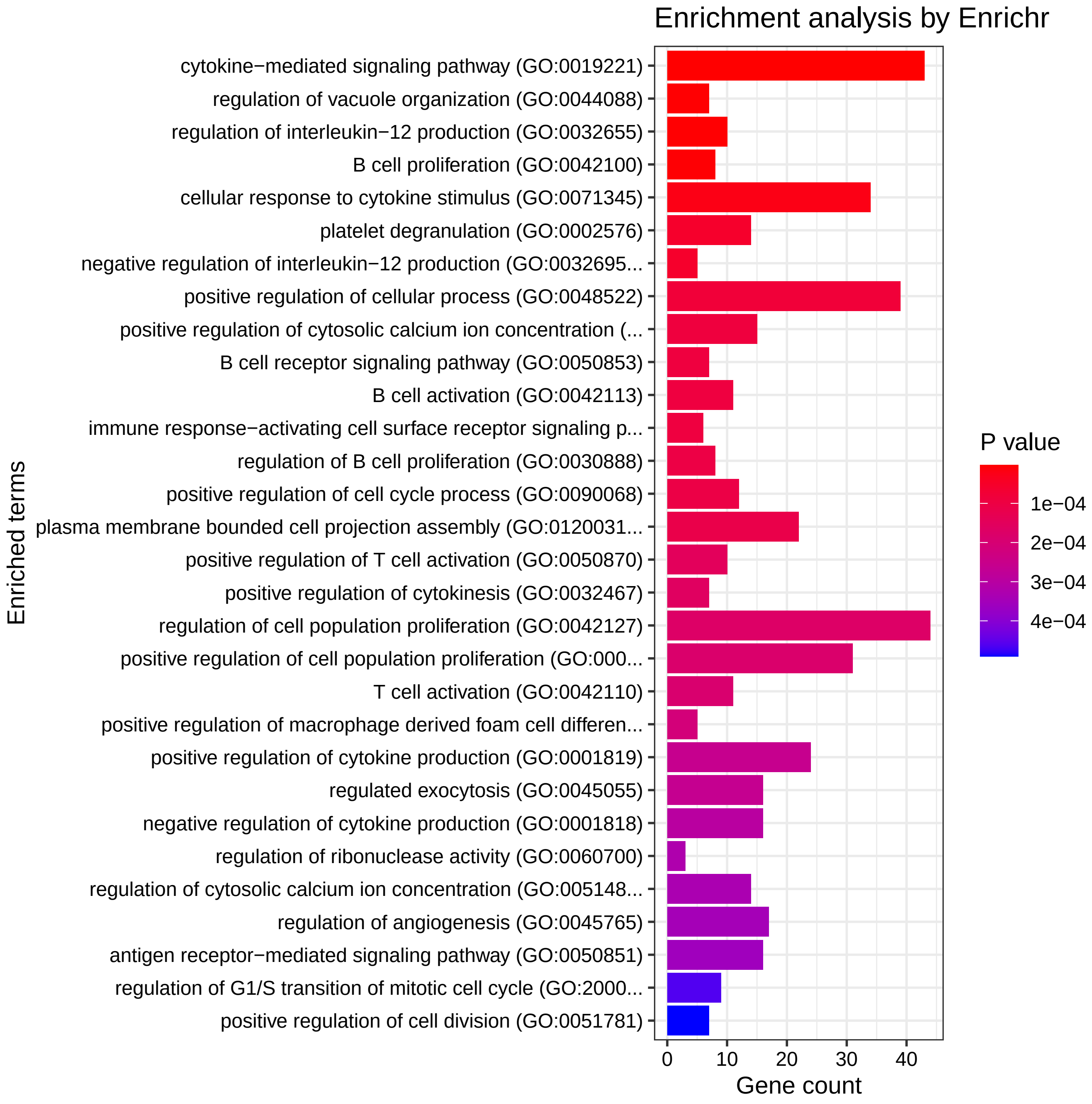

### Barplot_ReDisX_GSE59867_clus3_GO_CC.pdf

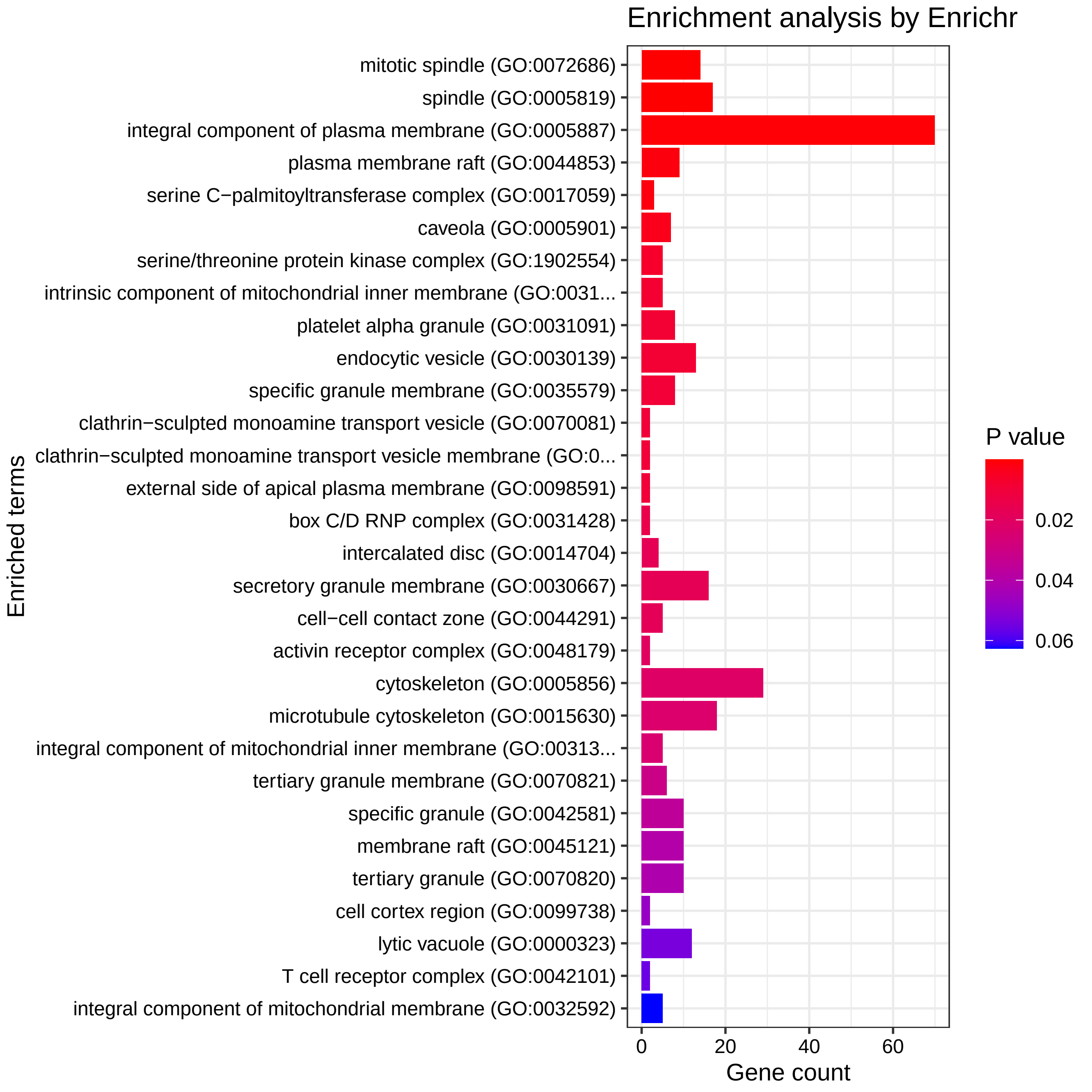

### Barplot_ReDisX_GSE59867_clus3_GO_MF.pdf

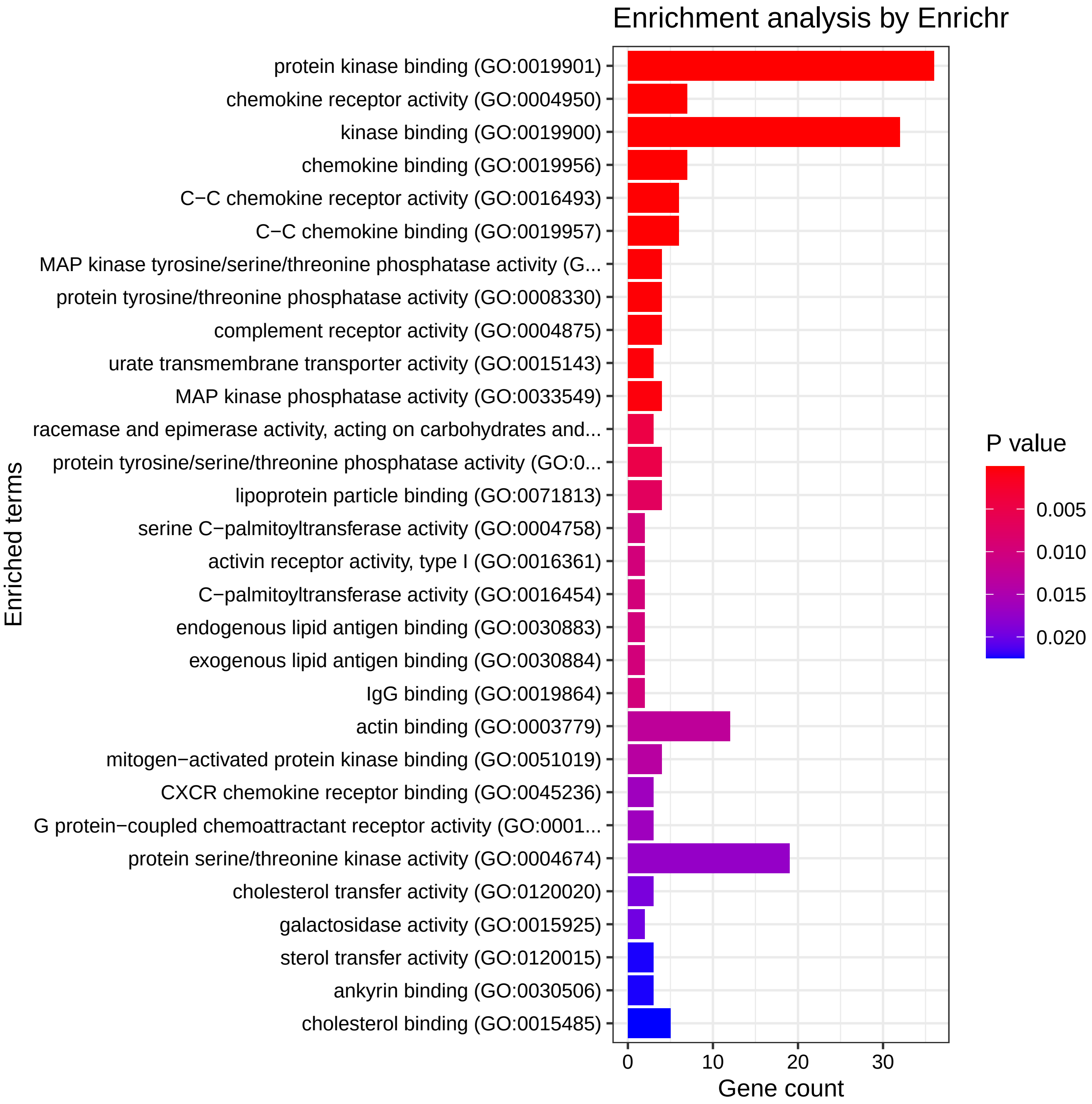

### Barplot_ReDisX_GSE59867_clus3_KEGG.pdf

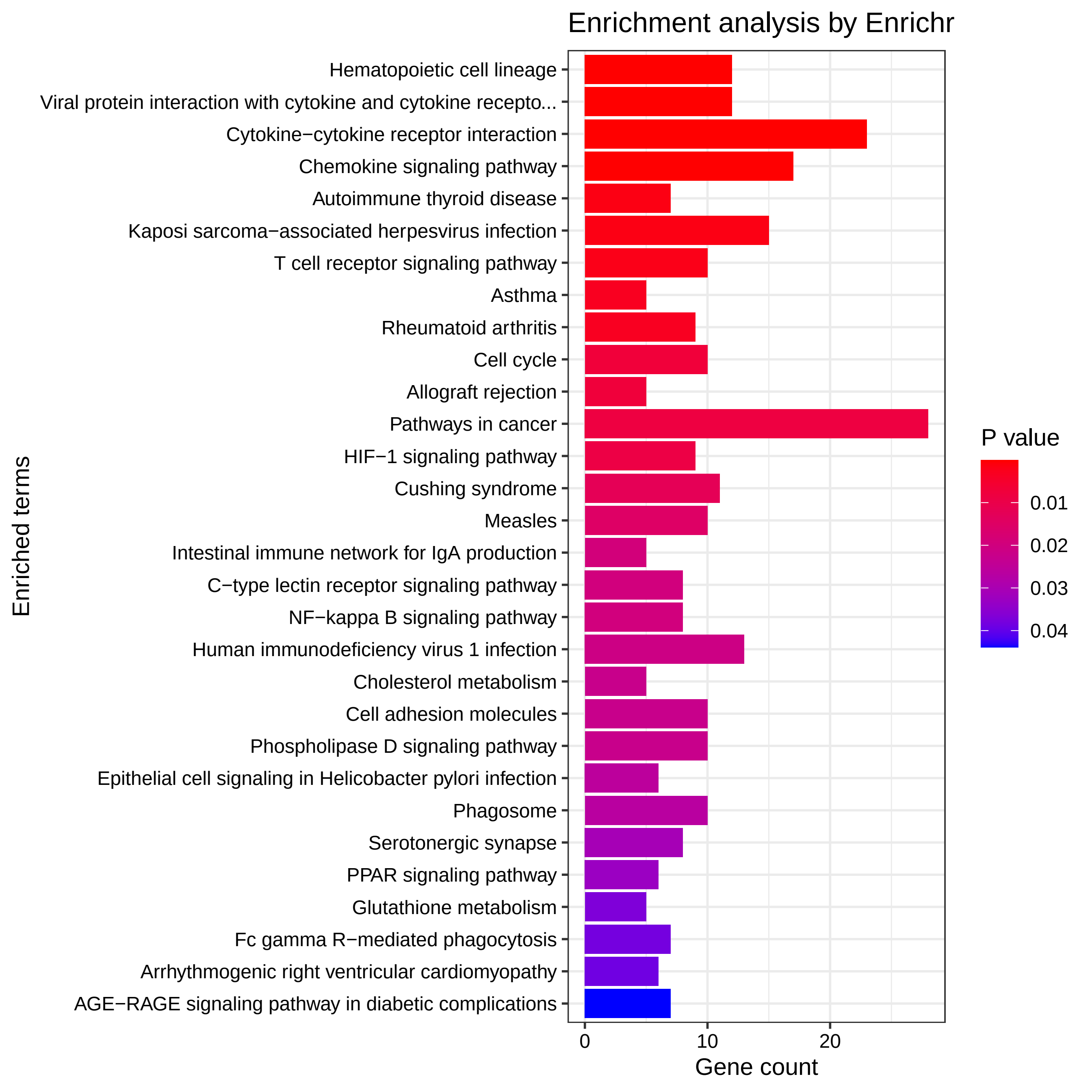

### clusterProfiler_gseDGN_Disgenet_GSE59867_GSE15573_ReDisXclus3_Dotplot.pdf

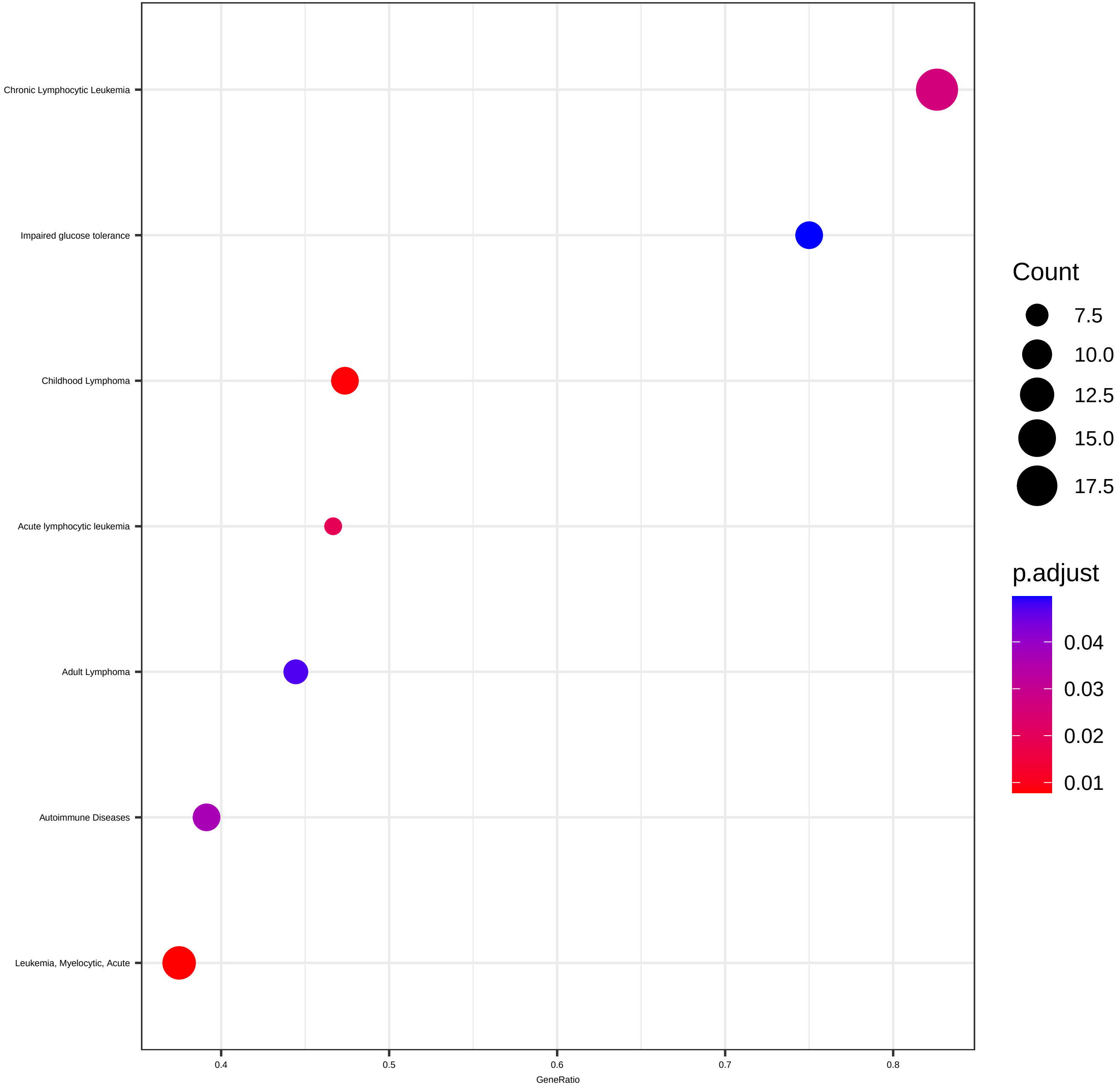

### clusterProfiler_gseKEGG_15573_59867_Dotplot.pdf

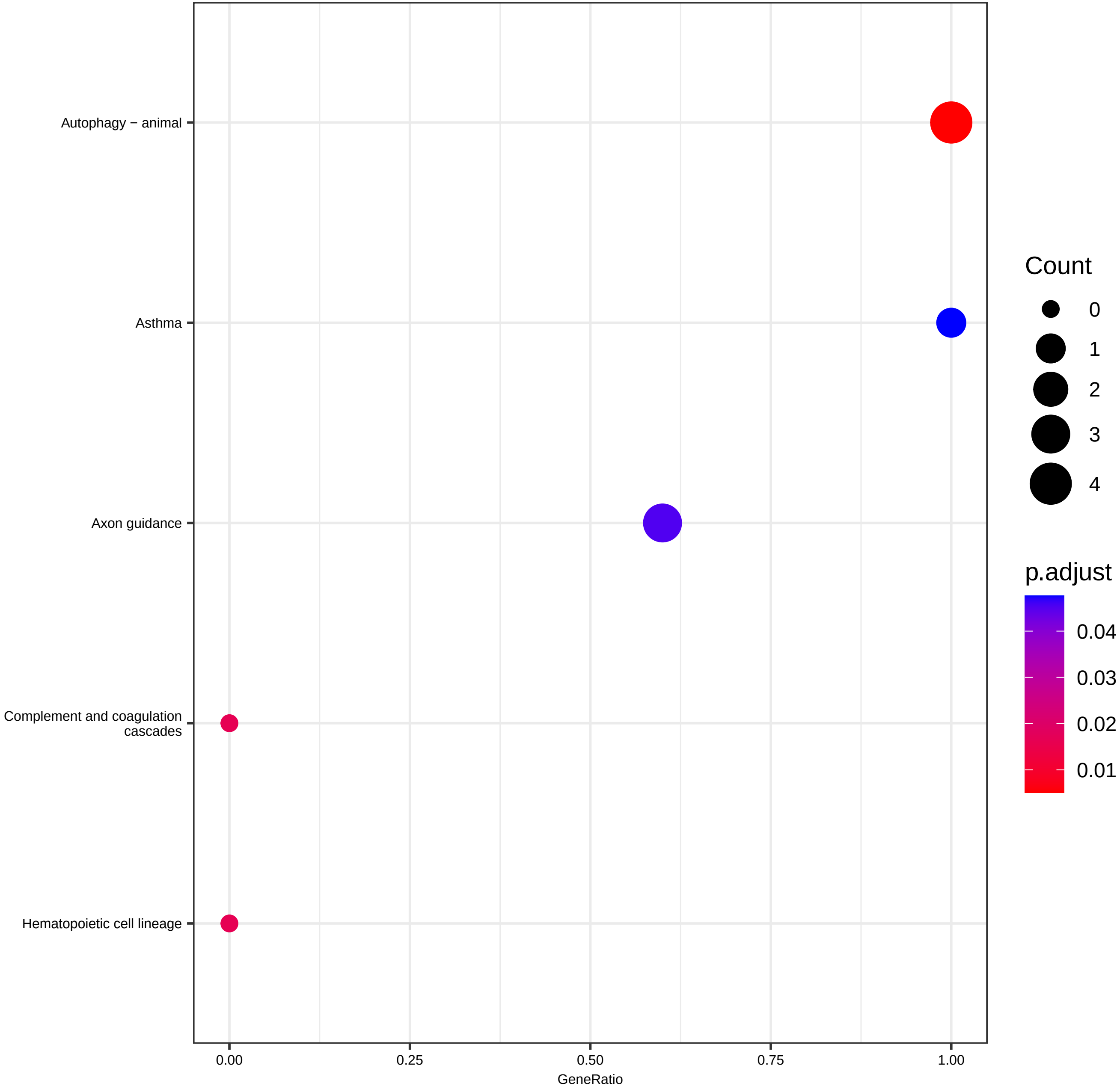

### Disgenet_GSE59867_ReDisXclus3_Dotplot.pdf

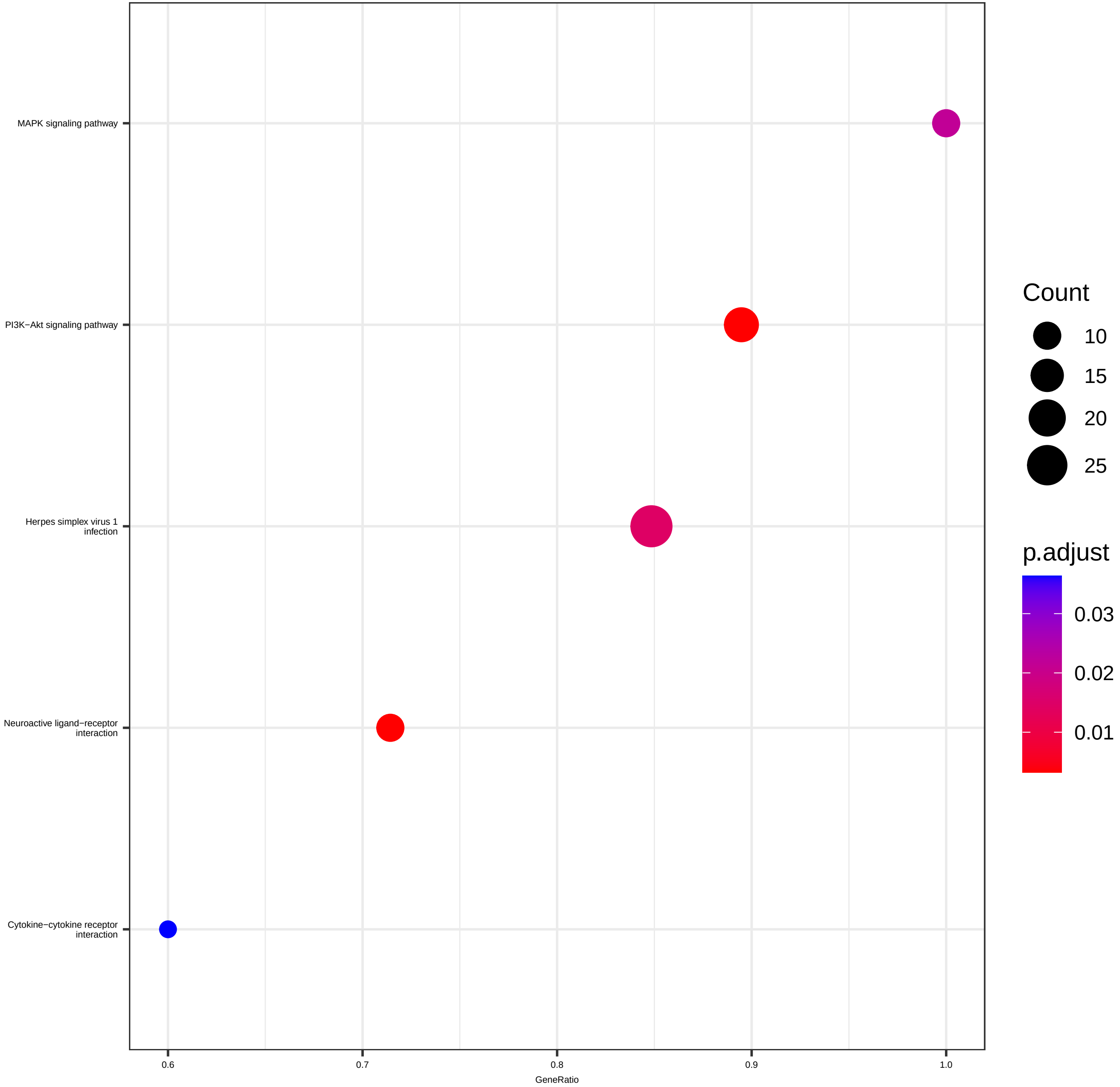

### GSEA_Disgenet_GSE59867_ReDisXclus3.pdf

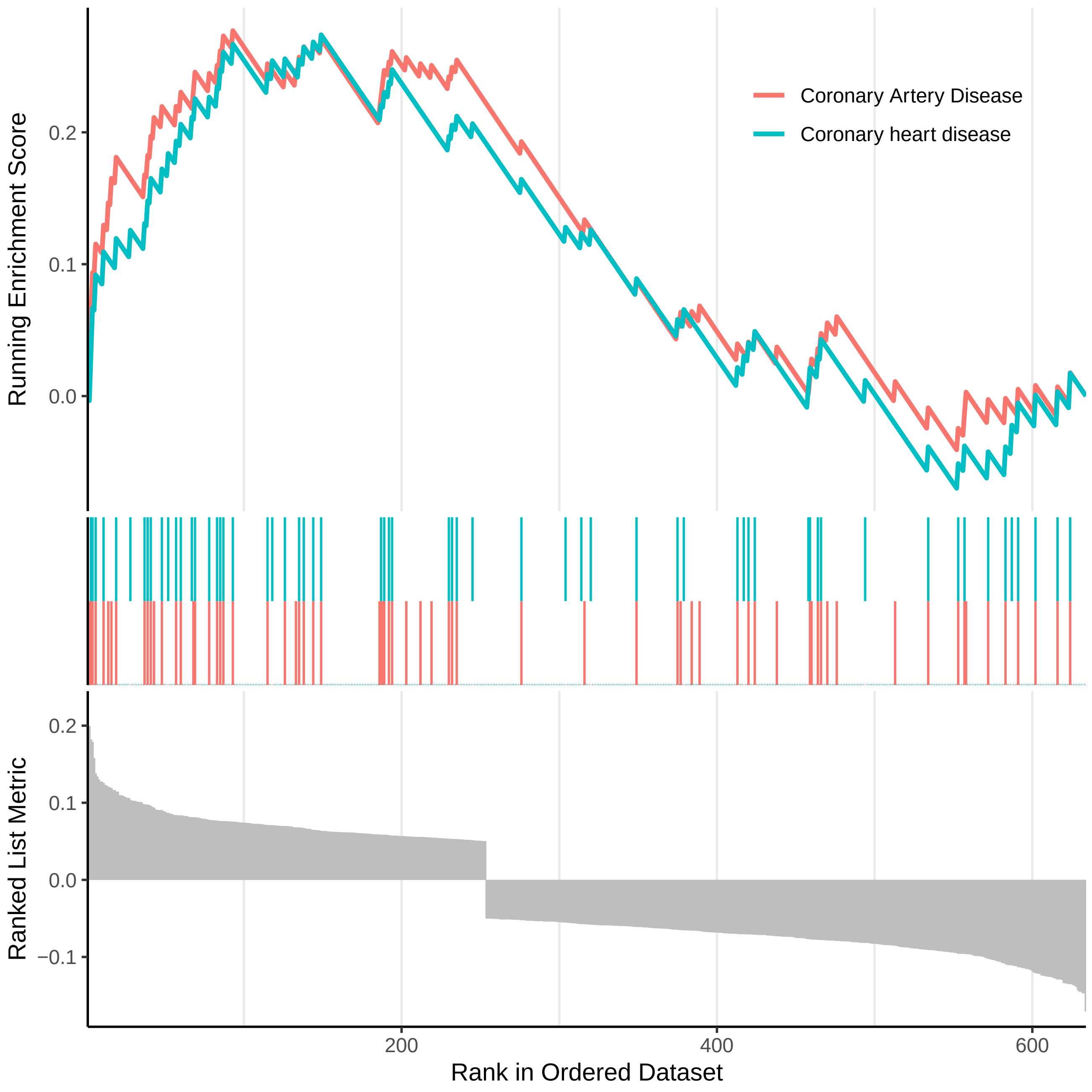
