## supplementary 3 for "ReDisX: a Continuous Max Flow-based framework to redefine the diagnosis of diseases based on identified patterns of genomic signatures": VENN_and_intersection.R

```
library(readr)
library(enrichR)
library(edgeR)
library(limma)
library(Glimma)
library(org.Hs.eg.db)
library(gplots)
library(RColorBrewer)
library(NMF)
library(pheatmap)
library(clusterProfiler)
library(ggplot2)
library(VennDiagram)
library(RColorBrewer)
library(dplyr)
library(grid)
library(org.Hs.eg.db)
library(enrichplot)
library(GOSemSim)
library(DOSE)


setEnrichrSite("Enrichr")
websiteLive <- TRUE
setwd("/home/yip/Documents/29_12_2021_ReDisX_analysis/result_v2/GSE77298_GSE93272_GSE59867_clus3")

ReDisX_GSE59867_clus3 <- read_csv("distict_gene_DisRedXclus3_GSE59867.csv")
ReDisX_GSE59867_logfold_change <- read_csv("ReDisX_DEG_toplist_CAD.csv")


GSE93272_RA <- read_csv("distict_gene_RA_vs_Control.csv")
GSE77298_RA <- read.csv("GSE77298_p0_05.csv",header=FALSE)
colnames(GSE77298_RA)="SYMBOL"

intersect_93272_59867=intersect(GSE93272_RA$SYMBOL,ReDisX_GSE59867_clus3$SYMBOL)
write.csv(intersect_93272_59867, 'GSE93272_intersect_GSE59867.csv')

intersect_93272_77298=intersect(GSE93272_RA$SYMBOL,GSE77298_RA$SYMBOL)
write.csv(intersect_93272_77298, 'GSE93272_intersect_GSE77298.csv')

intersect_77298_59867=intersect(GSE77298_RA$SYMBOL,ReDisX_GSE59867_clus3$SYMBOL)
write.csv(intersect_77298_59867, 'GSE77298_intersect_GSE59867.csv')


intersect_77298_59867_93272=intersect(intersect_77298_59867,GSE93272_RA$SYMBOL)
write.csv(intersect_77298_59867_93272, 'GSE77298_intersect_GSE59867_intersect_GSE93272.csv')


dbs <- c("GO_Molecular_Function_2021", "GO_Cellular_Component_2021", "GO_Biological_Process_2021","KEGG_2021_Human","DisGeNET")
if (websiteLive) {
  enriched <- enrichr(intersect_77298_59867, dbs)
}


pdf(file="enrich_GSE59867_GSE77298/Barplot_77298_59867_KEGG.pdf")
if (websiteLive) plotEnrich(enriched[[4]], showTerms = 30, numChar = 60, y = "Count", orderBy = "P.value")

dev.off()

pdf(file="enrich_GSE59867_GSE77298/Barplot_77298_59867_GO_MF.pdf")
if (websiteLive) plotEnrich(enriched[[1]], showTerms = 30, numChar = 60, y = "Count", orderBy = "P.value")

dev.off()
pdf(file="enrich_GSE59867_GSE77298/Barplot_77298_59867_GO_CC.pdf")
if (websiteLive) plotEnrich(enriched[[2]], showTerms = 30, numChar = 60, y = "Count", orderBy = "P.value")

dev.off()
pdf(file="enrich_GSE59867_GSE77298/Barplot_77298_59867_GO_BP.pdf")
if (websiteLive) plotEnrich(enriched[[3]], showTerms = 30, numChar = 60, y = "Count", orderBy = "P.value")

dev.off()
pdf(file="enrich_GSE59867_GSE77298/Barplot_77298_59867_DisGeNet.pdf")
if (websiteLive) plotEnrich(enriched[[5]], showTerms = 30, numChar = 60, y = "Count", orderBy = "P.value")

dev.off()

############# all _DEGs GSE59867_clus3

if (websiteLive) {
  enriched_ReDisX_GSE59867_clus3 <- enrichr(ReDisX_GSE59867_clus3$SYMBOL, dbs)
}


pdf(file="enrich_GSE59867_ReDisXclus3/Barplot_ReDisX_GSE59867_clus3_KEGG.pdf")
if (websiteLive) plotEnrich(enriched_ReDisX_GSE59867_clus3[[4]], showTerms = 30, numChar = 60, y = "Count", orderBy = "P.value")

dev.off()

pdf(file="enrich_GSE59867_ReDisXclus3/Barplot_ReDisX_GSE59867_clus3_GO_MF.pdf")
if (websiteLive) plotEnrich(enriched_ReDisX_GSE59867_clus3[[1]], showTerms = 30, numChar = 60, y = "Count", orderBy = "P.value")

dev.off()
pdf(file="enrich_GSE59867_ReDisXclus3/Barplot_ReDisX_GSE59867_clus3_GO_CC.pdf")
if (websiteLive) plotEnrich(enriched_ReDisX_GSE59867_clus3[[2]], showTerms = 30, numChar = 60, y = "Count", orderBy = "P.value")

dev.off()
pdf(file="enrich_GSE59867_ReDisXclus3/Barplot_ReDisX_GSE59867_clus3_GO_BP.pdf")
if (websiteLive) plotEnrich(enriched_ReDisX_GSE59867_clus3[[3]], showTerms = 30, numChar = 60, y = "Count", orderBy = "P.value")

dev.off()
pdf(file="enrich_GSE59867_ReDisXclus3/Barplot_ReDisX_GSE59867_clus3_DisGeNet.pdf")
if (websiteLive) plotEnrich(enriched_ReDisX_GSE59867_clus3[[5]], showTerms = 30, numChar = 60, y = "Count", orderBy = "P.value")

dev.off()


kegg_organism = "hsa"
KEGG_77298_59867 = enrichKEGG(gene= geneList_77298_59867,
                   organism     = kegg_organism,
                   minGSSize    = 1,
                   maxGSSize    = 20000,
                   pvalueCutoff = 0.05,
                   pAdjustMethod = "BH",
                   keyType       = "ncbi-geneid")


cluster_summary_KEGG_77298_59867<- data.frame(KEGG_77298_59867)
write.csv(cluster_summary_KEGG_77298_59867, "enrich_GSE59867_GSE77298/clusterProfiler_enrichKEGG_77298_59867.csv")


pdf(file="enrich_GSE59867_GSE77298/clusterProfiler_enriKEGG_77298_59867_Dotplot.pdf")

dotplot(KEGG_77298_59867 , showCategory=30,font.size = 6)
dev.off()

######################################################
####### GSEA

logfold_GSE_77298_59867=ReDisX_GSE59867_logfold_change[match(ReDisX_GSE59867_clus3$SYMBOL,ReDisX_GSE59867_logfold_change$SYMBOL,nomatch = 0),c(1,5)]
colnames(logfold_GSE_77298_59867)[1]="SYMBOL"


ids_77298_59867=bitr(logfold_GSE_77298_59867$SYMBOL, fromType = "SYMBOL", toType = "ENTREZID", OrgDb= org.Hs.eg.db)
dedup_ids_77298_59867 = ids_77298_59867[!duplicated(ids_77298_59867[c("SYMBOL")]),]

logfold_GSE_77298_59867["ENTREZID"]=dedup_ids_77298_59867$ENTREZID

logfold_GSE_77298_59867 = logfold_GSE_77298_59867[order(logfold_GSE_77298_59867$ReDisX_CAD_clus3_Control_DEG, decreasing = TRUE),]


geneList_77298_59867=logfold_GSE_77298_59867$ENTREZID
geneList_77298_59867=logfold_GSE_77298_59867$ReDisX_CAD_clus3_Control_DEG
names(geneList_77298_59867)=as.character(logfold_GSE_77298_59867$ENTREZID)


DisGenet_77298_59867  <- gseDGN(gene=geneList_77298_59867,
                       pvalueCutoff  = 0.05,
                       pAdjustMethod = "BH")
DisGenet_77298_59867  <- setReadable(DisGenet_77298_59867 , 'org.Hs.eg.db')

cluster_summary_DisGenet_77298_59867<- data.frame(DisGenet_77298_59867)
write.csv(cluster_summary_DisGenet_77298_59867, "enrich_GSE59867_ReDisXclus3/clusterProfiler_gseDGN_ReDisXclus3.csv")


pdf(file="enrich_GSE59867_ReDisXclus3/Disgenet_GSE59867_ReDisXclus3_Dotplot.pdf")
dotplot(DisGenet_77298_59867, showCategory=30,font.size = 4)
dev.off()
pdf(file="enrich_GSE59867_ReDisXclus3/GSEA_Disgenet_GSE59867_ReDisXclus3.pdf")
gseaplot2(DisGenet_77298_59867 , geneSetID = c(18,49))
dev.off()


##############################3 ReDisX_clus4 GSEA

ReDisX_GSE59867_clus4 <- read_csv("distict_gene_DisRedXclus4_GSE59867.csv")
logfold_clus4=ReDisX_GSE59867_logfold_change[match(ReDisX_GSE59867_clus4$SYMBOL,
                                                         ReDisX_GSE59867_logfold_change$SYMBOL,nomatch = 0),c(1,6)]
colnames(logfold_clus4)[1]="SYMBOL"


ids_clus4=bitr(logfold_clus4$SYMBOL, fromType = "SYMBOL", toType = "ENTREZID", OrgDb= org.Hs.eg.db)
dedup_ids_clus4 = ids_clus4[!duplicated(ids_clus4[c("SYMBOL")]),]

logfold_clus4["ENTREZID"]=dedup_ids_clus4$ENTREZID

logfold_clus4 = logfold_clus4[order(logfold_clus4$ReDisX_CAD_clus4_Control_DEG, decreasing = TRUE),]


#geneList_clus1=logfold_clus1$ENTREZID
geneList_clus4=logfold_clus4$ReDisX_CAD_clus4_Control_DEG
names(geneList_clus4)=as.character(logfold_clus4$ENTREZID)


DisGenet_clus4  <- gseDGN(gene=geneList_clus4,
                                pvalueCutoff  = 0.05,
                                pAdjustMethod = "BH")
DisGenet_clus4  <- setReadable(DisGenet_clus4 , 'org.Hs.eg.db')

cluster_summary_DisGenet_clus4<- data.frame(DisGenet_clus4)
write.csv(cluster_summary_DisGenet_clus4, "enrich_GSE59867_ReDisXclus4/clusterProfiler_DisGenet_clus4.csv")


pdf(file="enrich_GSE59867_ReDisXclus4/Disgenet_GSE59867_ReDisXclus4_Dotplot.pdf")
dotplot(DisGenet_clus4, showCategory=30,font.size = 4)
dev.off()
pdf(file="enrich_GSE59867_ReDisXclus4/GSEA_Disgenet_GSE59867_ReDisXclus4.pdf")
gseaplot2(DisGenet_clus4 , geneSetID = c(2,3,40))
dev.off()

kegg_organism = "hsa"
KEGG_59867_clus4 = gseKEGG(gene= geneList_clus4,
                              organism     = kegg_organism,
                              pvalueCutoff = 0.05,
                              pAdjustMethod = "BH")

cluster_summary_KEGG_59867_clus4 <- data.frame(KEGG_59867_clus4 )
write.csv(cluster_summary_KEGG_59867_clus4 , "enrich_GSE59867_ReDisXclus4/clusterProfiler_gseKEGG_59867_clus4 .csv")


pdf(file="enrich_GSE59867_ReDisXclus4/GSEA_KEGG_GSE59867_ReDisXclus4.pdf")
dotplot(KEGG_59867_clus4, showCategory=30,font.size = 4)
dev.off()

############# all _DEGs GSE59867_clus4

if (websiteLive) {
  enriched_ReDisX_GSE59867_clus4 <- enrichr(ReDisX_GSE59867_clus4$SYMBOL, dbs)
}


pdf(file="enrich_GSE59867_ReDisXclus4/Barplot_ReDisX_GSE59867_clus4_KEGG.pdf")
if (websiteLive) plotEnrich(enriched_ReDisX_GSE59867_clus4[[4]], showTerms = 30, numChar = 60, y = "Count", orderBy = "P.value")

dev.off()

pdf(file="enrich_GSE59867_ReDisXclus4/Barplot_ReDisX_GSE59867_clus4_GO_MF.pdf")
if (websiteLive) plotEnrich(enriched_ReDisX_GSE59867_clus4[[1]], showTerms = 30, numChar = 60, y = "Count", orderBy = "P.value")

dev.off()
pdf(file="enrich_GSE59867_ReDisXclus4/Barplot_ReDisX_GSE59867_clus4_GO_CC.pdf")
if (websiteLive) plotEnrich(enriched_ReDisX_GSE59867_clus4[[2]], showTerms = 30, numChar = 60, y = "Count", orderBy = "P.value")

dev.off()
pdf(file="enrich_GSE59867_ReDisXclus4/Barplot_ReDisX_GSE59867_clus4_GO_BP.pdf")
if (websiteLive) plotEnrich(enriched_ReDisX_GSE59867_clus4[[3]], showTerms = 30, numChar = 60, y = "Count", orderBy = "P.value")

dev.off()
pdf(file="enrich_GSE59867_ReDisXclus4/Barplot_ReDisX_GSE59867_clus4_DisGeNet.pdf")
if (websiteLive) plotEnrich(enriched_ReDisX_GSE59867_clus4[[5]], showTerms = 30, numChar = 60, y = "Count", orderBy = "P.value")

dev.off()

ReDisX_GSE59867_clus4_DisGenet=data.frame(DisGenet_clus4@result)

ReDisX_GSE59867_clus4_DisGenet_RA=ReDisX_GSE59867_clus4_DisGenet[c("C0003873"),]
ReDisX_GSE59867_clus4_DisGenet_RA_core=ReDisX_GSE59867_clus4_DisGenet_RA["core_enrichment"]
```
